# Engineering circuits of human iPSC-derived neurons and rat primary glia

**DOI:** 10.1101/2022.11.07.515431

**Authors:** Sophie Girardin, Stephan J. Ihle, Arianna Menghini, Magdalena Krubner, Leonardo Tognola, Jens Duru, Tobias Ruff, Isabelle Fruh, Matthias Müller, János Vörös

**Affiliations:** Laboratory of Biosensors and Bioelectronics, Institute for Biomedical Engineering, Department of Electrical Engineering and Information Technology, University and ETH Zürich, Zürich, Switzerland; Chemical Biology & Therapeutics, Novartis Institutes for BioMedical Research, Basel, Switzerland

**Keywords:** bottom-up neuroscience, iPSC-derived neurons, glial cells, in vitro neural circuits, microelectrode arrays, magnesium, drug testing, spheroids, neural engineering

## Abstract

Novel *in vitro* platforms based on human neurons are needed to improve early drug testing and address the stalling drug discovery in neurological disorders. Topologically controlled circuits of human induced pluripotent stem cell (iPSC)-derived neurons have the potential to become such a testing system. In this work, we build *in vitro* co-cultured circuits of human iPSC-derived neurons and rat primary glial cells using microfabricated polydimethylsiloxane (PDMS) structures on microelectrode arrays (MEAs). Such circuits are created by seeding either dissociated cells or pre-aggregated spheroids at different neuron-to-glia ratios. Furthermore, an antifouling coating is developed to prevent axonal overgrowth in undesired locations of the microstructure. We assess the electrophysiological properties of different types of circuits over more than 50 days, including their stimulation-induced neural activity. Finally, we demonstrate the effect of magnesium chloride on the electrical activity of our iPSC circuits as a proof-of-concept for screening of neuroactive compounds.

## 1 INTRODUCTION

Despite the increasing incidence of central nervous system (CNS) diseases worldwide, drug discovery targeting brain-related diseases is stalling (Howes and Mehta, 2021). CNS drug candidates often show promising results in all stages of preclinical tests, but fail in human clinical trials (Kesselheim et al., 2015). The low success rate in clinical trials shows that current preclinical CNS drug testing systems are not suitable to efficiently screen for new candidate molecules. Animal models, which are widely used in preclinical tests, often lack predictive value of efficacy in humans and fail to recapitulate relevant CNS disease phenotypes (Gribkoff and Kaczmarek, 2017). There is a clear need for the development of versatile *in vitro* test platforms incorporating relevant, disease-specific CNS cells, along with a suitable functional readout to identify effective drug candidates. A potential approach to build such a drug testing platform is to use the technical tools recently developed in the field of “bottom-up” neuroscience, in combination with disease-relevant cell types such as human induced pluripotent stem cell (iPSC)-derived neurons. Bottom-up neuroscience focuses on building and studying elementary circuits of neurons to infer the principles of more complex assemblies, with the main goal of getting a better understanding of the mechanisms behind information processing in the brain (Aebersold et al., 2016). Bottom-up neuroscience tools allow to build multiple, topologically-controlled circuits of tens to hundreds of neurons, which can be electrically stimulated and recorded from in parallel using microelectrode arrays (MEA) (Forro et al., 2018; Ihle et al., 2022; Duru et al., 2022; Girardin et al., 2022).

Besides the more and more refined technological tools to engineer *in vitro* neural networks, recent advances in the field of human iPSCs are another promising step towards improving drug testing systems (Xu and Zhong, 2013; Little et al., 2019; Farkhondeh et al., 2019). iPSC-derived neurons are a scalable source of genetically relevant cells for *in vitro* models. They can potentially replace the cells of animal origin, which are often poor predictors of efficacy and adverse side effects in humans. Networks of randomly connected iPSC-derived neurons have also been grown on MEAs to characterize their electrical activity (reviewed in Keller and Frega (2019)), for example in neurotoxicity testing (Kasteel and Westerink, 2017; Tukker et al., 2018) and to study neurodegenerative diseases (Mossink et al., 2021).

Several CNS drug screening platforms based on iPSC-derived neurons have been reported (Ahfeldt et al., 2017), but they usually rely exclusively on imaging assays. For example, two platforms based on random iPSC-derived neurons could successfully be used to identify Alzheimer’s disease drug candidates, using an imaging-based quantification of tau aggregation (Medda et al., 2016) and of amyloid beta (Kondo et al., 2017). A few platforms tailored to other neurological diseases have been reported, such as the use of iPSC-derived dopaminergic neurons for Parkinson’s disease drug screening (Moreno et al., 2015) or the use of iPSC-derived motoneurons with the potential to be used to study amyotrophic lateral sclerosis (Mo et al., 2020; Wainger et al., 2014). Fantuzzo et al. (2020) developed a platform based on a 96-well plate, where each well is divided into two compartments. Compartments were seeded with excitatory and inhibitory iPSC-derived neurons. Electrical activity was screened using calcium imaging. While calcium imaging can provide relevant information for functional screenings, an advantage of MEAs is the possibility to electrically stimulate the neuronal cultures, giving access to information-rich readouts. MEAs can also record the extracellular neural activity of a culture at multiple locations with higher temporal resolution. MEAs are particularly suitable for drug discovery, because repeated measurements can be performed on the same array over time, for example to measure the effect of adding a pharmacological molecule on the spontaneous and stimulation-induced electrical activity of a cultured network (Johnstone et al., 2010).

A reliable, fast, and efficient method to differentiate human iPSCs into neurons is through the overexpression of the gene neurogenin-2 (NGN2) as first reported by Zhang et al. (2013). Human iPSC-derived neurons obtained through the overexpression of NGN2, termed “induced” neurons or iNeurons, mostly exhibit properties typical of excitatory cortical neurons. However, compared to primary neurons of animal origin, human iPSC-derived neurons often suffer from poor survival after differentiation (Chen et al., 2021). In spite of this challenge, human NGN2 iNeurons have been successfully used with bottom-up neuroscience tools to build *in vitro* circuits composed of very few cells (Girardin et al., 2022). Human NGN2 iNeurons are routinely co-cultured with mice or rat primary astrocytes to support their maturation and function (Zhang et al., 2013; Frega et al., 2017; Fernandopulle et al., 2018; Rhee et al., 2019).

Astrocytes have received a lot of scientific attention in the past two decades thanks to a still growing body of evidence on the central roles that this glial cell type plays in brain development and function (Chandrasekaran et al., 2016). In addition to ensuring homeostasis and providing metabolic support to neurons, astrocytes are also involved in the facilitation of synapse formation (Ullian et al., 2001), the regulation of synaptic transmission (Fields and Stevens-Graham, 2002; Dallérac et al., 2013), and the clearance of neurotransmitters from the synaptic cleft (Weber and Barros, 2015). Dysfunctions of astrocytes contribute to the initiation and the propagation of many neurological diseases, while healthy astrocytes can help disease recovery (Verkhratsky et al., 2012). Besides astrocytes, primary glial cell preparations often contain microglia, another type of glial cells (Uliasz et al., 2012). As the resident macrophages of the CNS, microglia play a key role in CNS disorders (Perry et al., 2010). Because of their prominent role in CNS diseases, we decided to integrate rat primary glial cells to our *in vitro* system and test their effect on human iNeurons.

Here we present the use of a versatile platform to build multiple small circuits of different ratios of human iNeurons and rat primary glial cells, initially seeded as either dissociated cells or spheroids. We compare cultures of dissociated cells to cells aggregated into spheroids, as three-dimensional (3D) culture systems have received a lot of interest for drug testing (Jensen and Teng, 2020). Spheroids consist of glial cells and neurons pre-assembled into spherical co-cultures using commercially available microwells (Cvetkovic et al., 2018). We investigate the electrophysiological properties of such circuits, in particular their stimulation-evoked activity for more than 50 days in *vitro* (DIV). To show the potential of such a platform for drug testing, we evaluate the effect of the sequential addition of magnesium chloride to the cell medium. Our approach is highly versatile and provides the readout of neural activity with high signal-to-noise-ratio from 15 circuits in parallel. Furthermore, it is possible to measure the response to electrical stimulation over weeks or even months. We believe that such a system has the potential to be used for screening pharmacological compounds in the early stages of drug discovery.

## 2 MATERIALS AND METHODS

### 2.1 PDMS microstructures

Polydimethylsiloxane (PDMS) microstructures were based on a previously published design and prepared as described in Girardin et al. (2022). PDMS microstructures consist of a micropatterned two-layer 120 μm thin membrane fabricated by Wunderlichips (Switzerland). One microstructure contains an array of 5×3 4-node circuits as shown in Fig. 1A. Each node has a cylindrical hole with a diameter of 170 μm and is connected to the neighboring nodes through 4 μm high microchannels.

**Figure 1.**
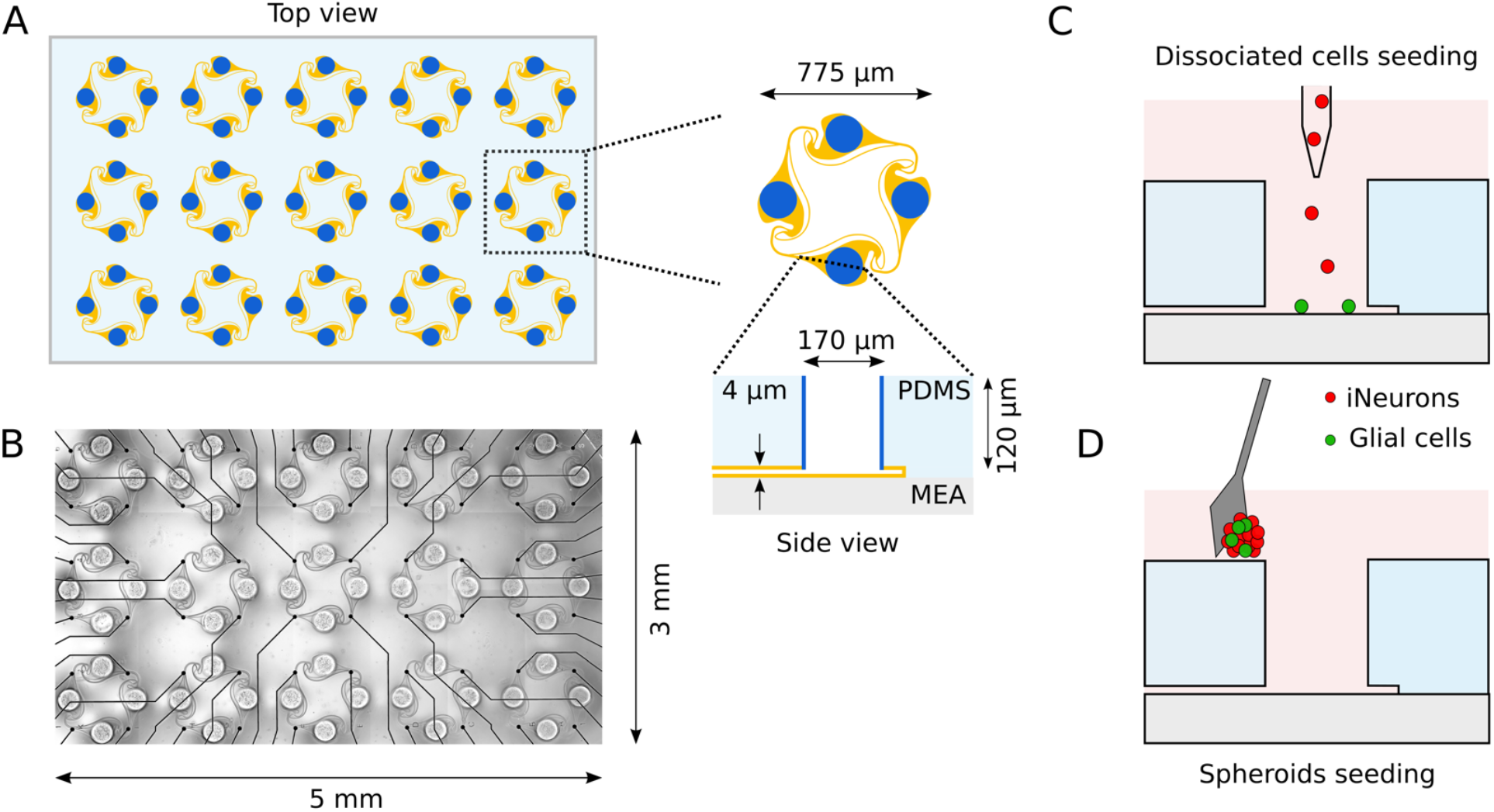
Overview of the *in vitro* platform used to build small circuits of iNeurons and glial cells. **(A)** Schematic top view of a typical PDMS microstructure used in this work. The PDMS microstructure is a 120 μm thick membrane containing 15 circuits. A circuit consists of four nodes (blue) connected by microchannels (orange). Nodes are cylindrical holes with a diameter of 170 μm. Microchannels are 4 μm-high to prevent soma from migrating into them, while still allowing axons to grow in them. **(B)** The microchannels of the PDMS microstructure can be aligned on top of a 60-electrode MEA, allowing to record from and stimulate the bundle of axons passing on top of the electrodes. **(C)** Two different seeding modalities were used in the present work: dissociated cells (C) or spheroids (D). The type of cells seeded were either iNeurons (red) or a mixture of iNeurons and glial cells (green). To seed dissociated cells, glial cells were first pipetted into the nodes, followed by iNeurons. **(D)** Cells were left in non-adhesive microwells for two days to form spheroids, were then individually placed in the nodes by pushing them into the wells with a micro-knife. *Schematics are not to scale.*

#### 2.1.1 PDMS coating

To prevent axons from growing on the top surface of the PDMS microstructure, a coating based on the antifouling poly(vinylpyrrolidone) (PVP) was applied on top of the PDMS. Perfluorophenyl azide (PFPA) was grafted to a poly(allylamine) (PAAm) backbone at a ratio of 1:4 and used as an adhesion layer to covalently bind PVP to the top surface of the PDMS. A solution of 3 mg/mL of PAAm-grafted-PFPA (PAAm-g-PFPA) was synthesized as detailed in Weydert et al. (2019). Briefly, PFPA-NHS (ATFB-NHS, RL-2045, Iris Biotech) dissolved in ethanol was added to PAAm (283215, Sigma Aldrich) dissolved in a potassium carbonate buffer (pH 8.5, 791776, Sigma Aldrich). Prior to use, the PAAm-g-PFPA was diluted to 0.1 mg/mL in 2:3 HEPES1/ethanol mixture. HEPES1 consists of 10 mM HEPES at pH 7.4, obtained by diluting 0.05 M hydroxyethyl piperazine ethanesulfonic acid (HEPES) (94717, Sigma Aldrich) in deionized water.

PDMS microstructures were placed on a sacrificial glass slide and plasma cleaned for 2 min (18 W PDC-32G, Harrick Plasma). A droplet of about 10 μL containing 0.1 mg/mL PAAm-g-PFPA was added on top of each PDMS microstructure to cover the entire top surface of the microstructure. The microstructures were left at room temperature and protected from light for 30 min in a humid environment to prevent the solution from evaporating. The PDMS microstructures were then rinsed with 2:3 HEPES1/ethanol and with ultrapure water (Milli-Q, Merck-MilliPore). PVP (1.3 MDa, 437190, Sigma Aldrich) was dissolved in ethanol to a concentration of 10 mg/mL. A droplet of 10 μL was added on top of the PDMS microstructures (already coated with the PFPA adhesion layer) and gently blow dried with nitrogen. The coated PDMS was exposed to 254 nm light for 5 min, then soaked in methanol for 1 hr, where the methanol solution was replaced every 15 min. Finally, the PDMS microstructures were ultrasonicated for 5 min in fresh methanol, rinsed in ultrapure water and left in ultrapure water until use. The coated PDMS microstructures were used within two days.

### 2.2 Substrate preparation

Staining experiments and coating trials were performed on 35-mm diameter glass bottom dishes (HBST-3522T, WillCo Wells). Electrophysiological experiments were performed using 60-electrode microelectrode arrays (60MEA500/30iR-Ti-gr, Multi Channel Systems). Substrates were prepared according to the protocol detailed in Girardin et al. (2022), which includes a Poly-D-lysine (PDL, P6407, Sigma Aldrich) coating of substrates and alignment of the PDMS microstructures on the substrates (MEA or glass-bottom dish), as shown in Fig. 1B. For MEAs, a single PFPA-PVP coated PDMS microstructure was used and aligned, whereas for glass-bottom dishes, four PFPA-PVP PDMS microstructures were used, without the need for alignment. Substrates were then stored at 4°C and used for cell seeding within two days.

### 2.3 Cell culture

#### 2.3.1 iPSC differentiation

Exactly as also described in Girardin et al. (2022), human iPSCs were generated following a previously published protocol (Giorgetti et al., 2019) and transfected with a doxycycline-inducible Neurogenin-2 (NGN2) gene. Differentiation into neurons was induced by a 3-day exposure to doxycycline as reported in Russell et al. (2018). Differentiated iNeurons were then cryogenized as aliquots of 1 · 10^6^ to 8 · 10^6^ cells in heat inactivated fetal bovine serum (FBS, 10270-106, Thermo Fisher) containing 5% v/v dimethyl sulfoxide (DMSO). Cryogenized aliquots of iNeurons were kindly provided by Novartis and stored in liquid nitrogen until use.

#### 2.3.2 Generation of RFP NGN2 line

To generate the red fluorescent protein (RFP)-positive NGN2 line, RFP was cloned into a PiggyBac (PB) plasmid (Russell et al., 2018) under the CAG promoter. This plasmid was nucleofected into the NGN2 iPS line (obtained as described above) using a nucleofection kit (Human Stem Cell Nucleofector Kit 1, VPH-5012, Lonza) containing 4 μg of RFP PB construct and 1 μg of dual helper plasmid. The nucleofection was performed using an Amaxa Nucleofector II (Lonza Bioscience, program B-016). After selection with 1 μg/ml puromycin, clones were picked and analyzed for homogeneous fluorescence.

#### 2.3.3 Cell culture medium preparation

Glia medium consisted of DMEM (61965-026) with 10% v/v FBS and 1% v/v antibiotic-antimycotic 100X (15240-062, Thermo Fisher) and was used for cultures of glial cells only. The culture medium used for iNeuron or iNeuron-glia cultures was Neurobasal Differentiation (NBD) medium. NBD was prepared from Neurobasal medium (21203-049) by adding 1% v/v GlutaMAX (35050-061) and 1% v/v Pen-Strep (15070-063), all from Thermo Fisher. 1 mL of B27 supplement (17504-044, Thermo Fisher), 0.5 mL of N2 supplement (17502-048, Thermo Fisher), 50 μL of brain-derived neurotrophic factor (BDNF, 10 μg/mL, 450-10, PeproTech) and 50 μL of glial-derived neutrophic factor (GDNF, 10 μg/mL, 450-02, PeproTech) were added freshly to 50 mL of Neurobasal medium.

In the first 24 hr after thawing, iNeurons were cultured in NBD+RI, *i.e.* NBD supplemented with 10 μM of Rho-Kinase Inhibitor Y27632 (10 mM in PBS, 688000, Sigma Aldrich), as this was shown to improve survival of thawed iPSC-derived cells (Chen et al., 2021). From day in *vitro* (DIV) 0 to 7, NBD was also supplemented with 5 μg/mL of laminin (11243217001, Sigma Aldrich) to help with neuron survival (Girardin et al., 2022). From DIV 4 to 9, glia-containing samples were cultured with NBD supplemented with 2 μM of cytosine-β-D-arabinofuranoside (AraC, 500 μM in ultrapure water, C1768, Sigma Aldrich) to inhibit astrocyte and microglia proliferation. From DIV 10 onwards, NBD was supplemented with 2.5% FBS (NBD+FBS) to support astrocyte survival (Frega et al., 2017).

#### 2.3.4 Primary rat glial cell cultures

Primary rat glial cultures were obtained from cortices of E18 embryos of pregnant Sprague-Dawley rats (EPIC, ETH Phenomics Center, Switzerland). Animal experiments were approved by the Cantonal Veterinary Office Zurich, Switzerland. The cortices were dissociated following a previously published protocol (Ihle et al., 2022). Glial cells were then seeded at a density of 100k cells/mL into T25 flasks (90026, TPP Techno Plastic Products AG), which had previously been coated with 0.1 mg/mL of PDL in PBS for 2 hr at 37°C. The medium was exchanged after 2 days to get rid of dead cells, after which a complete change to 5 mL of fresh glia medium was performed every four days. 10 to 14 days after the initial seeding, once the culture reached confluency, it was split using trypsin/0.25% EDTA (25200-056, Thermo Fisher) and reseeded into two non-coated T25 flasks.

Glial cells were passaged once or twice before being used in PDMS microstructures. For seeding into PDMS microstructures or for the generation of spheroids, glial cell flasks were first rinsed with warm PBS, then left in 3 mL of trypsin/0.25% EDTA at 37°C for 10 to 15 min. They were then centrifuged for 5 min at 100 g, resuspended into fresh NBD and counted using an automatized cell counter (Cell Countess, Invitrogen).

#### 2.3.5 Dissociated sample preparation: seeding glia and iNeurons into PDMS microstructures

Three different ratios of iNeurons-to-glia were used to make samples: iNeurons only, 5:1 iNeurons-glia and 2:1 iNeurons-glia. In order to be able to compare samples, the number of iNeurons was kept constant at 90k cells/cm^2^.

Prior to cell seeding, the PBS in the PDMS-containing substrates was replaced with fresh NBD and placed at 37°C for at least 1 hr in a CO_2_ controlled environment. For glia-containing samples, glial cells were detached from their flask and seeded at a density of 18k cells/cm^2^ (5:1 samples) and 45k cells/cm^2^ (2:1 samples). Glial cells were then left in the samples for at least 4 hr before seeding the iNeurons.

A frozen aliquot of iNeurons was taken out of liquid nitrogen and rapidly thawed at 37°C. The 1 mL thawed cell solution was transferred dropwise into 4 mL of warm NBD and centrifuged for 5 min at 200 g. Cells were resuspended in NBD+RI at a concentration of 1 mio cells/mL. The cell solution was passed through a 40 μm strainer (CSS013040, BioFilJet) and counted. iNeurons were seeded on the samples at a concentration of 90k cells/cm^2^ (Fig. 1C), and were resuspended with a P1000 pipette at least twice in the first hour after seeding. A complete medium change was done about 1 hr after seeding to remove floating dead cells, where the NBD+RI medium was replaced with NBD+RI+laminin.

#### 2.3.6 Spheroid sample preparation

##### 2.3.6.1 Spheroid generation

A commercially available microwell culture plate was used to generate spheroids (AggreWell 400 24-well plate, 34411, StemCell Technologies) following the manufacturer’s instructions. The microwells were first coated with an anti-adherence rinsing solution (07010, StemCell Technologies) and centrifuged at 2,000 g for 5 min. The solution was then replaced with fresh NBD+RI+laminin and left at 37°C until use. Glial cells were detached from their flask and iNeurons were thawed as detailed above. iNeurons were seeded to form spheroids of 250 cells on average (relying on the statistical distribution of the cells in the microwell). Glial cells were added to two thirds of the iNeuron-containing microwells, at a ratio of either 1:5 or 1:2. The microwells were then placed in an incubator for two days.

##### 2.3.6.2 Seeding spheroids into PDMS microstructures

After two days in the microwells, spheroids of iNeurons, 5:1 iNeurons-glia and 2:1 iNeurons-glia were seeded into the nodes of PDMS microstructures. They were first taken out of the microwells with a P100 pipette, pipetting about 50-80 spheroids at a time. The spheroids were then placed in a small Petri dish and a P10 pipette was used to aspirate 60 spheroids under a stereo microscope. The spheroids were then slowly dispensed on top of the PDMS. An ethanol-disinfected dissection micro-knife (10316-14, Fine Science Tools) was used to gently push individual spheroids into the nodes of the PDMS microstructure (Fig. 1D) under a stereo microscope. Once all 60 nodes of the PDMS structure were filled with a spheroid, the micro-knife was used to gently push them to the bottom of the node to ensure that they did not float out of the PDMS node when moving the sample. Samples were kept for up to 15 min outside of the incubator. If this time was not sufficient to seed all spheroids, the sample was placed back into the incubator and the seeding process was continued 15 min later.

#### 2.3.7 Culture maintenance

Approximately half of the medium was exchanged two to three times a week, using the medium detailed in Section “Cell culture medium preparation”. For dissociated samples, DIV 0 was considered to be the day of seeding iNeurons into the PDMS microstructures. For spheroid samples, DIV 0 was considered as the day of seeding iNeurons into the spheroid-forming microwells.

### 2.4 Staining and imaging

#### 2.4.1 Immunofluorescence staining

Samples were immunostained following the protocol reported in Girardin et al. (2022). After 4 % paraformaldehyde (1.00496, Sigma Aldrich) fixation, 0.2 % Triton X-100 (X100, Sigma Aldrich) permeabilization and blocking of non-specific binding with 0.2 % Triton-X and 3 % bovine serum albumin (BSA, A7906, Sigma Aldrich), samples were incubated overnight at 4 °C with a solution of PBS containing 0.2 % Triton-X, 3 % BSA and the primary antibodies. Primary antibodies were chicken anti-GFAP (1:1000, AB4674, Abcam), rabbit anti-S100β(AB34686, Abcam), and mouse anti-MAP2 (1:250, 13-1500, Thermo Fisher). On the next day, samples were rinsed with PBS and incubated with a secondary antibody solution in PBS for 2 h at room temperature. Secondary antibodies were goat anti-mouse DyLight 405 (1:800, 35500BID, Thermo Fisher), goat anti-chicken Alexa Fluor 488 (1:800, A11039, Thermo Fisher) and donkey anti-rabbit Alex Fluor Plus 647 (1:800, A32795, Thermo Fisher). Depending on the experiment, Hoechst 33342 (H3570, Thermo Fisher) was added to the antibody solution at a working concentration of 2 μM instead of the DyLight 405 antibody. The samples were rinsed three times with PBS and left in PBS for imaging.

#### 2.4.2 Image acquisition

Microscopy images were acquired using a confocal laser scanning microscope (FluoView 3000, Olympus) with a 20X objective (NA 0.5, UPLFLN20XPH, Olympus). Four channels were acquired: 405 nm (Hoechst or DyLight 405), 488 nm (Alexa Fluor 488), and 647 nm (Alexa Fluor Plus 647) in addition to phase contrast brightfield images.

#### 2.4.3 Image analysis

Microscopy images were processed and analyzed using the open-source software Fiji (Schindelin et al., 2012). Because stained somas are bigger and thus much brighter than stained axons on microscopy images, the intensity of the axons was enhanced by using a pixel logarithm operator. This was applied to all the representative fluorescent images shown in the figures of this paper, except for the immunofluorescent stainings shown in Fig. 2 and 3.

**Figure 2.**
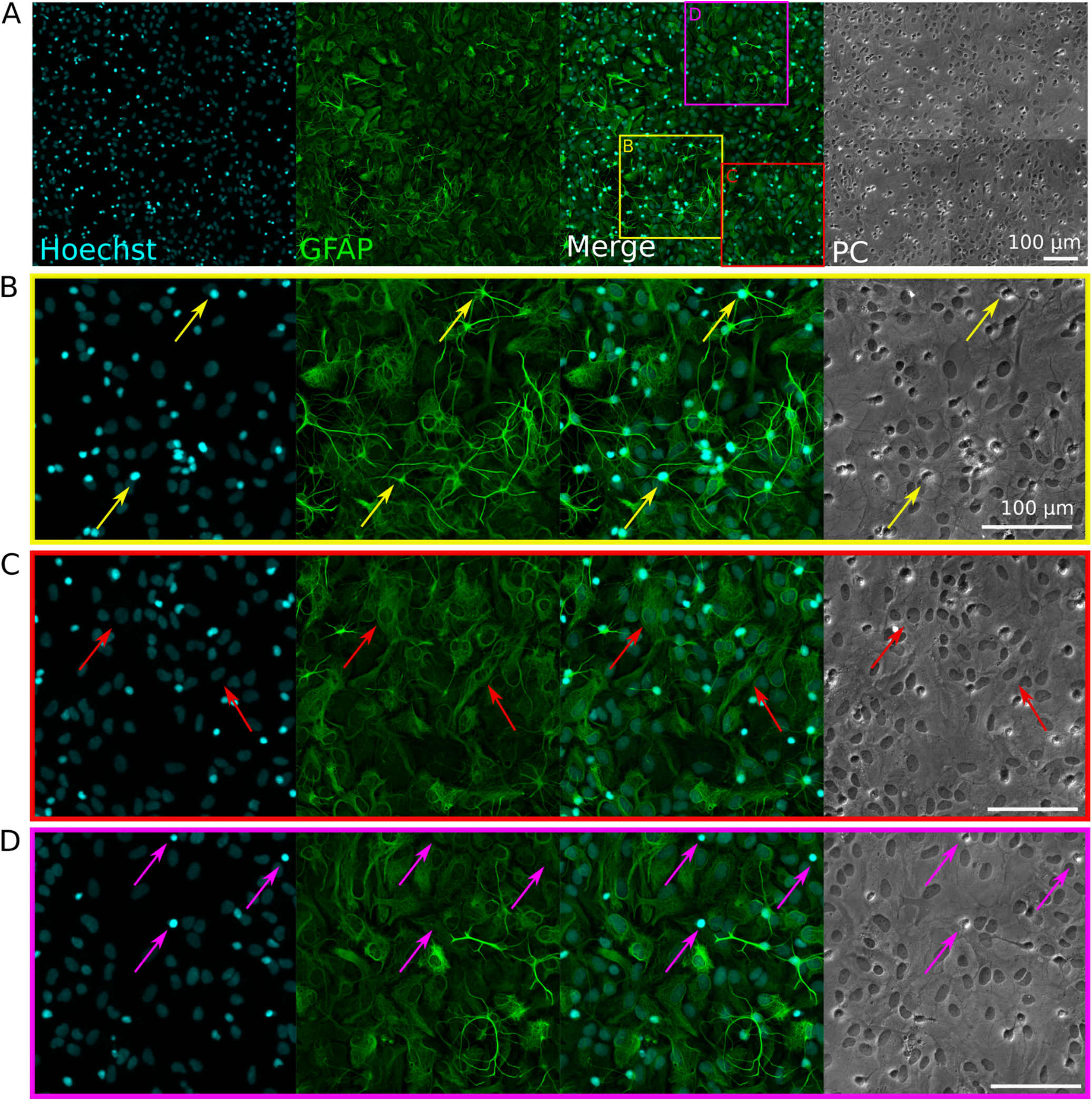
Immunofluorescence staining of rat primary glial cell cultures. Hoechst (cyan) stains nuclei, GFAP (green) stains astrocytes. ‘PC’: phase contrast (brightfield) images. **(A)** Representative image of the immunostained glia culture. The three colored squares visible on the ‘Merged’ image correspond to the zoom-in images shown in B-D. **(B)** The yellow arrows show examples of astrocytes with a “star-shape” morphology and a small, bright, and round nucleus. **(C)** The red arrows show examples of astrocytes with a “flat” morphology and a bigger, dim, and oval nucleus. **(D)** The pink arrows show examples of microglia with small, bright nuclei, which appear as bright spots in the phase-contrast image.

**Figure 3.**
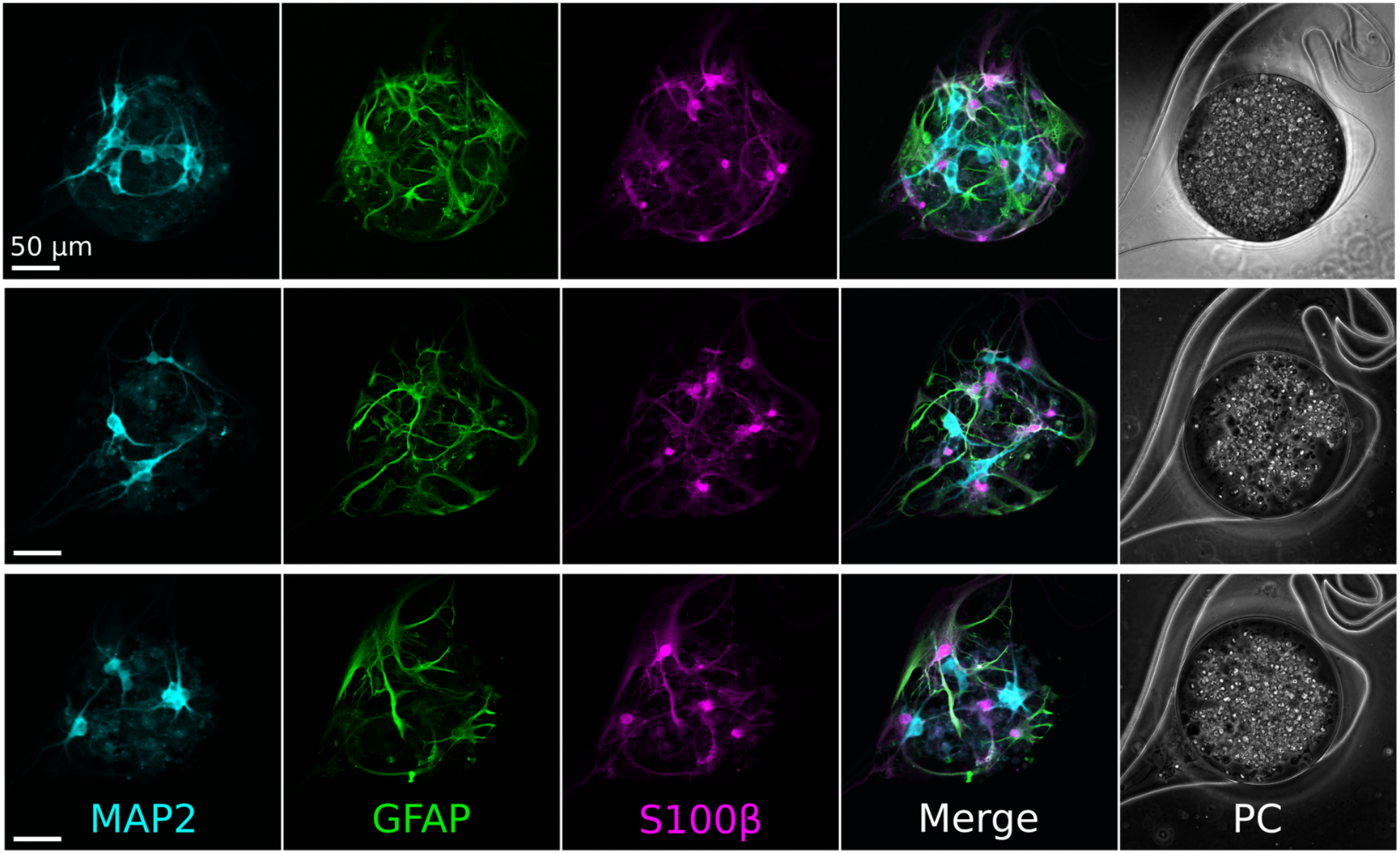
Immunofluorescence staining of the cells contained in the nodes of an iNeurons/glia circuit at DIV 16. Three representative examples are shown. The neuronal marker MAP2 (cyan) stained the iNeurons, while the astrocyte markers GFAP (green) and S100β (pink) stained the rat primary astrocytes. ‘PC’: phase contrast images

##### 2.4.3.1 PDMS overgrowth analysis

To determine if the PFPA-PVP coating effectively prevented axons from growing on top of the PDMS, three glass-bottom dishes (each containing four PDMS microstructures) with non-coated PDMS microstructures and three glass-bottom dishes with PFPA-PVP coated PDMS microstructures were prepared. All 6 samples were seeded with 5:1 red fluorescent protein (RFP)-expressing iNeurons-glia. Images of the top surface of the 60 nodes were acquired at DIV 7, 14, 28, and 35. These were cropped into individual circuits and manually inspected to determine whether or not axons were present.

##### 2.4.3.2 Visual assessment of the independence of the circuits

At DIV 50, all MEA samples used in the electrophysiology tests were imaged. The images were cropped into individual circuits and manually inspected to determine if axons grew outside of the intended area, as well as the number of empty nodes in each circuit.

### 2.5 Electrophysiology

Six different types of circuits were tested: dissociated iNeurons, dissociated 5:1 iNeurons-glia, dissociated 2:1 iNeurons-glia, and spheroids containing 250 iNeurons on average without any glial cells as well as at a iNeuron-glia ratio of 5:1 and 2:1. Each condition was repeated three times, *i.e.* on three different MEAs containing 15 circuits each (45 circuits per condition).

#### 2.5.1 Data acquisition

During electrophysiology recording sessions, each MEA was taken out of the incubator and placed in a MEA headstage recording unit (MEA2100-Systems, Multi Channel Systems), heated to 37 °C using the internal headstage heating place and a temperature controller (TCO2, Multi Channel Systems), and kept at 5% CO_2_ (Pecon #0506.00). The MEA was placed in the headstage 5 min before starting the recording session to acclimate to the environment. Data were acquired at 20 kHz from all 60 electrodes.

#### 2.5.2 Spike detection

Raw data were band-passed filtered with a 300 Hz high-pass filter (Butterworth, 2nd order). The baseline noise of the signal was characterized for each electrode using the median absolute deviation (MAD)(Quiroga et al., 2004). Spikes were detected by identifying positive signal peaks above a threshold of 7 times the filtered baseline noise. Successive events within 2 ms were discarded to avoid multiple detection of the same spike.

#### 2.5.3 Spontaneous electrical activity

##### 2.5.3.1 Mean firing rate and percentage of active electrodes

Electrophysiological activity was assessed over 8 weeks from 5-min recordings of spontaneous activity of a MEA. Each electrode’s mean firing rate (MFR) was calculated as the number of spikes detected per electrode divided by the recording time. An electrode was considered active if its MFR was above 0.1 Hz. Only active electrodes were used for subsequent analysis.

##### 2.5.3.2 Spike train directionality

To characterize the directionality of pairs of consecutive spikes, the timestamps of the detected spikes of all four electrodes of each circuit were inspected. Two spikes were considered to be related to each other if they fired within 5 ms of each other. The pre-spike and post-spike electrode was extracted for each pair of consecutive spikes. The frequency of each pair of pre-/post-electrodes was calculated and used to assess the overall directionality of consecutive spike pairs. Only pairs of spikes taking place on two adjacent electrodes (1-2, 2-3, 3-4 and 4-1) were taken into account for the calculation of the overall directionality of spike trains. Pairs of spikes taking place twice on the same electrode or on diagonally opposed electrodes (1-3 or 2-4) were discarded, as they cannot be linked to either clockwise or counter-clockwise directionality. The inactive electrodes of a circuit were ignored. The percentage of clockwise spike trains of each circuit was averaged over all conditions to give the total percentage of clockwise circuits.

#### 2.5.4 Stimulation-induced electrical activity

To investigate the response of a circuit to a repeated electrical stimulus, we recorded the response of all four electrodes of a circuit upon stimulation of one of its electrodes. The stimulus was a 400 μs biphasic square pulse with amplitude of 500 mV (first positive then negative). This stimulus was sequentially applied to each electrode of a circuit (top left, top right, bottom right, bottom left) at 4 Hz for 2 min. One set of stimuli thus corresponded to 480 repeats of the stimulus on the same electrode. An idle time of 30 sec was added between each set of stimuli.

The number of spikes induced in a circuit by the stimulation of one of its electrodes was assessed by summing the number of spikes detected on all four electrodes of the circuit in the 250 ms following the stimulus during the 2 min of stimulation of the electrode. An electrode was considered to be “activity-inducing” if at least 480 spikes were detected during the set of stimuli, *i.e.* if each electrical stimulus elicited at least one spike on average.

#### 2.5.5 Effect of magnesium on spontaneous and stimulation-induced electrical activity

To investigate the effect of magnesium on the MFR and on the stimulation-induced activity, concentrated magnesium chloride (1 M in H_2_O, 63069, Sigma Aldrich) was sequentially added to the medium of different MEAs. Recording sessions consisted of 5 min of spontaneous activity, 20 min of stimulation (alternating 2 min of stimulation per electrode) and another 5 min of spontaneous activity recording. The initial magnesium ion (Mg^2+^) concentration in NBD was 0.81 mM. The concentrated magnesium chloride solution was used to sequentially increase the concentration of extracellular Mg^2+^ by 2.5, 5, 7.5, and 10 mM. Before starting the addition of magnesium chloride, the cell medium was first switched to 1.1 mL of fresh, warm NBD+FBS. The MEA was left to equilibrate in the incubator for at least 40 min before starting the first recording session. After the first recording session, 2.75 μL of aqueous 1 M magnesium chloride were added to the medium to increase the Mg^2+^ concentration by 2.5 mM, bringing the extracellular concentration to 3.31 mM. The MEA was left in the incubator for about 40 min before repeating the measurements. After having added an overall 10 mM of magnesium chloride and performed the corresponding recordings, the medium was fully exchanged, bringing the extracellular Mg^2+^ concentration back to 0.81 mM. The sample was left to equilibrate for 40 min, before performing a last set of recordings.

## 3 RESULTS

In this work, we report the building of circuits made of human iPSC-derived NGN2 neurons (iNeurons) and rat primary glial cells. The iNeurons/glia circuits are characterized using imaging, electrophysiology recordings and electrical stimulation. A proof-of-concept of their potential use in drug screening is presented by assessing the effect of elevated extracellular magnesium concentrations on the electrical activity of these circuits.

The circuits are topologically constrained by PDMS microstructures and consist of four nodes containing the cell bodies, connected together with microchannels which are high enough for neurites to enter, but too low for soma to migrate into (Fig. 1A). The PDMS microstructures are based on a previously published design that ensures predominantly clockwise axon growth between the nodes (Forro et al., 2018; Girardin et al., 2022). The PDMS microstructures contain an array of 15 circuits and can be placed on a 60-electrode MEA, where the microchannels are aligned on the electrodes (Fig. 1B). Cells are seeded into the microstructure either as dissociated cells (Fig. 1C) or as pre-aggregated spheroids (Fig. 1D). Once axons have grown in the microchannels, they can be stimulated and their extracellular electrical activity can be recorded with the microelectrodes.

### 3.1 Immunofluorescent characterization of glial cells and iNeurons

#### 3.1.1 Rat primary glial cells

Cultures of primary glial cells were established from E18 rat cortices. Such cultures are typically mostly composed of astrocytes, but can also contain other glial cell types, such as microglia (Uliasz et al., 2012). Importantly, the primary glia cultures had to be neuron-free to avoid seeding rat neurons in the PDMS microstructures along with the glial cells. To verify this, we performed an immunofluorescent staining on a confluent culture of glial cells passaged once, using antibodies against MAP2 (a neuronal marker) and GFAP (an astrocyte marker), as well as the nucleus marker Hoechst. The results of the staining are visible in Fig. 2.

None of the cells were stained by the anti-MAP2 antibody (see Fig. S1), confirming the absence of neurons. The anti-GFAP stain shows that astrocytes presented two distinct types of morphologies: star-shaped and brightly GFAP-stained astrocytes (Fig. 2B, yellow arrows), or flat and dimly GFAP-stained astrocytes (Fig. 2C, red arrows). These two morphologies can also be distinguished in the phase-contrast images, with the edges of the star-shaped astrocytes appearing brighter and the nuclei of the flat astrocytes standing out as dark spots. In the Hoechst phase, the morphology of the nuclei of the two types of astrocytes is also clearly recognizable, with a rounder, smaller, and brighter nucleus for the star-shaped astrocytes, and a bigger, dimmer, and more oval nucleus for flat astrocytes. Some bright nuclei without a surrounding GFAP-positive region were also visible in the culture (Fig. 2D, pink arrows). Such cells appeared as bright, round spots in the phase-contrast images, which is the typical morphology of microglia on top of an astrocyte layer in glial cultures (Georgieva et al., 2018). Overall, these results indicate that the glial cells consist mostly of astrocytes, with a subset of microglia, and that they can be used in co-culture with human iNeurons without the risk of introducing rat neurons.

#### 3.1.2 Circuits of iNeurons and rat glia

We sequentially seeded rat glial cells and human iNeurons in PDMS microstructures to confirm that they could be used to build small circuits. We successfully obtained circuits composed of a mixture of the two cell types. To verify the identity of the iNeurons and of the astrocytes, immunofluorescent staining was performed on iNeuron/glial circuits at 15 days *in vitro* (DIV). The antibodies were targeted to MAP2 (marking the neurons), GFAP and S100β (both astrocyte markers). Immunostaining results are presented in Fig. 3. iNeurons stained positively for MAP2, but not for GFAP or S100β, indicating that the neuronal differentiation did not get impaired by the close proximity to astrocytes. Most astrocytes expressed both GFAP and S100β, but some preferentially expressed one or the other protein. GFAP is a cytoskeleton protein and its antibody is expected to label the extensive branching of astrocytes, while S100β is a cytoplasmic calcium-binding peptide whose antibody labels the cell bodies of small astrocytes with less extensive branching (Jurga et al., 2021). In contrast to the staining performed on proliferating cultures of astrocytes, astrocytes inside of the structures mostly presented a star-like morphology.

In all samples, a lot of dead iNeurons are visible in the phase-contrast images. These are mostly iNeurons which recovered poorly from the post-cryopreservation thawing process and died during the first week of culture. iNeuron survival at low densities in PDMS microstructures was extensively discussed in a previous work, where it was shown that despite the presence of dead cells, circuits of iNeurons can survive over months, form synapses and have stable neural activity (Girardin et al., 2022). We did not investigate iNeuron survival again in the present work. Since the iNeurons-to-glia ratios “5:1” and “2:1” refer to the initially seeded ratios, the true iNeurons-to-glia ratio can deviate over time due to difference in viability of iNeurons and glial cells over the life cycle of a culture.

### 3.2 An antifouling coating is necessary to prevent axons from growing on top of the PDMS

We built circuits of human iNeurons and rat glial cells in PDMS microstructures at different ratios and with two different seeding modalities (as dissociated cells or as spheroids) and labeled them with a live-dead and nuclei stain (Fig. S2). While they could successfully be built, cultured over weeks and imaged, these circuits presented a major flaw: in addition to growing inside of the microchannels, axons also grew along the vertical walls of the PDMS nodes and onto the top surface of the PDMS microstructures. As a result, in less than a week, most of the 15 circuits of a microstructure ceased to be physically separated. This is not desirable, as it goes against the efforts of constraining the topology and compartmentalization of the neuronal circuits.

To prevent neurites from growing on top of the PDMS, the top surface of the microstructure can be coated with a neuron-repellent molecule. iNeurons can survive for months in PDMS microstructures. As we are interested in recording their electrical activity over several weeks, it is necessary to use a coating that is resistant over similar time spans. We thus selected the antifouling molecule PVP, which can be attached to the PDMS using a PAAm-*g*-PFPA adhesion layer, a surface functionalization method that was reported to be stable over several weeks when used in cultures of rat primary neurons (Weydert et al., 2019). The long-term adhesion of the coating relies on the UV-sensitive molecule PFPA, an aromatic compound with an azide (N_3_) functional group. Upon exposure to UV-C light (254 nm), the azide group reacts with carbon double bonds, carbon rings, methyl groups, or amine groups, effectively forming covalent bonds with any organic molecule containing one of these groups (Liu and Yan, 2010).

We thus used PAAm-g-PFPA/PVP (referred to as “PFPA-PVP” from now on) to coat the top surface of the PDMS microstructures. The PDMS was coated while located on a sacrificial glass slide, so that the bottom surface of the PDMS did not get functionalized and could still reliably adhere to the coverslip or MEA glass surface after coating. We compared the growth of axons on top of 12 PFPA-PVP coated PDMS microstructures and 12 non-coated PDMS microstructures. The samples were seeded with red fluorescent protein-expressing iNeurons (RFP iNeurons) and glial cells at an initial ratio of 5:1. Fig. 4A and B show the top surface of a non-coated and of a PFPA-PVP coated PDMS microstructure after DIV 35, with the network of neurites visible in red. Such fluorescent images were taken every week for five weeks and used to assess the presence of axon growth on the top surface of the PDMS for each circuit (Fig. 4C). In the non-coated microstructures, 80% of the circuits had axons growing on top at DIV 7 and 100% of them had axons on top at DIV 35. The slight decrease in axon overgrowth in non-coated samples between DIV 7 and DIV 13 is due to the fact that in the early days, the network of axons at the top of the PDMS is fragile and medium exchange is sometimes sufficient to disturb some of the axons.

**Figure 4.**
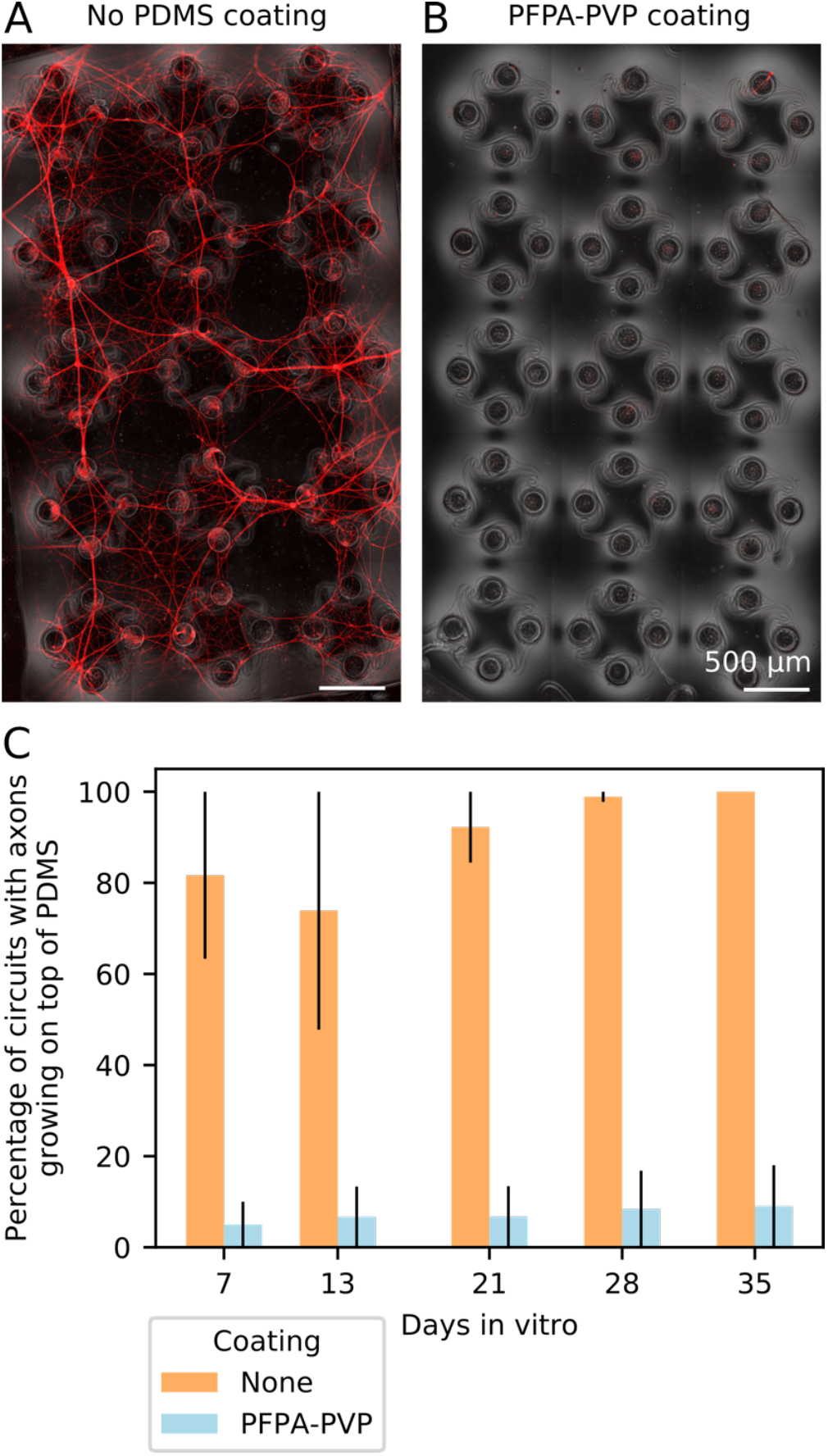
Coating the top surface of the PDMS microstructure prevents axons from growing there. **(A)** Representative image of the top surface of a non-coated PDMS microstructure at DIV 35. Axons from RFP-expressing iNeurons grow from the nodes and connect neighboring circuits together, disrupting the compartmentalization of the cells into individual circuits. **(B)** Representative image of the top surface of a PDMS microstructure coated with PFPA-PVP at DIV 35. The coating prevents axons from exiting the nodes and connecting the different circuits. **(C)** Quantification of the percentage of circuits where axons could grow out of the node and onto the top surface of the PDMS microstructure for both non-coated and PFPA-PVP coated PDMS. For each bar, N = 12 PDMS microstructures of 15 circuits each. Error bars show the SEM.

The PFPA-PVP coating effectively prevented 90% of the growth at all timepoints, with a slight decrease of performance over time. The few circuits of PFPA-PVP-coated microstructures which showed axon overgrowth were always located next to the edge of the structure, with axons from iNeurons growing from outside of the PDMS (Fig. S3). These were probably due to a non-perfect coating on the edges and to a rather thin microstructure edge (< 100 μm). For the MEA experiments presented below, slightly thicker microstructures were used (120 μm) and care was given to properly coat also the edge of the microstructures. No axon growth was observed on top of the PDMS microstructures on any of these MEA samples.

### 3.3 Building different types of circuits of iNeurons and glial cells on microelectrode arrays

Having found an effective coating to prevent axonal growth on top of the PDMS, we then built iNeuron/glia circuits on MEAs to investigate their electrical activity. Six different types of circuits were built using three different iNeuron-to-glia seeding ratios (1:0, 2:1, and 5:1) and two different seeding modalities (dissociated and spheroids). The seeding ratio of neurons-to-glia of 2:1 was selected as it roughly corresponds to the ratio that is present in the cortex (von Bartheld et al., 2016). Since only part of the initially seeded iNeurons survive, the ratio of 5:1 was picked as an intermediary point of comparison (16.7% of glia instead of 33.3% of glia). Three MEAs of each of the six different types of circuits were prepared for a total of 45 circuits per condition. RPF-expressing iNeurons were used to allow for regular live imaging without the need to repetitively add a live cell stain.

#### 3.3.1 Verifying the absence of electrically active cells in the glial cell cultures

To ensure that the primary glial cells did not have any detectable level of electrical activity, we seeded 3 MEAs with dissociated glial cells and 3 MEAs with spheroids of glial cells and recorded their spontaneous electrical activity after 7, 14, and 21 DIV. No action potentials were detected in the recordings (data not shown), confirming that all electrical activity recorded on the neuron-containing samples can be attributed solely to the iNeurons.

#### 3.3.2 Effect of glial cells on the morphology of iNeuron circuits

The growth and morphology of circuits were investigated by imaging in the first few days of culture, as well as after 2.5 and 7 weeks (DIV 17 and 50). Representative images of one circuit of each dissociated condition are shown in Fig. 5 and of one circuit of each spheroid condition is shown in Fig. 6, with both the RFP-expressing neurons (red) and the phase contrast (greyscale) images. For the three spheroid conditions, cells were first assembled in microwells for two days before being transferred into the PDMS microstructures. Therefore, an example image of the spheroids in the microwells is shown before DIV2.

**Figure 5.**
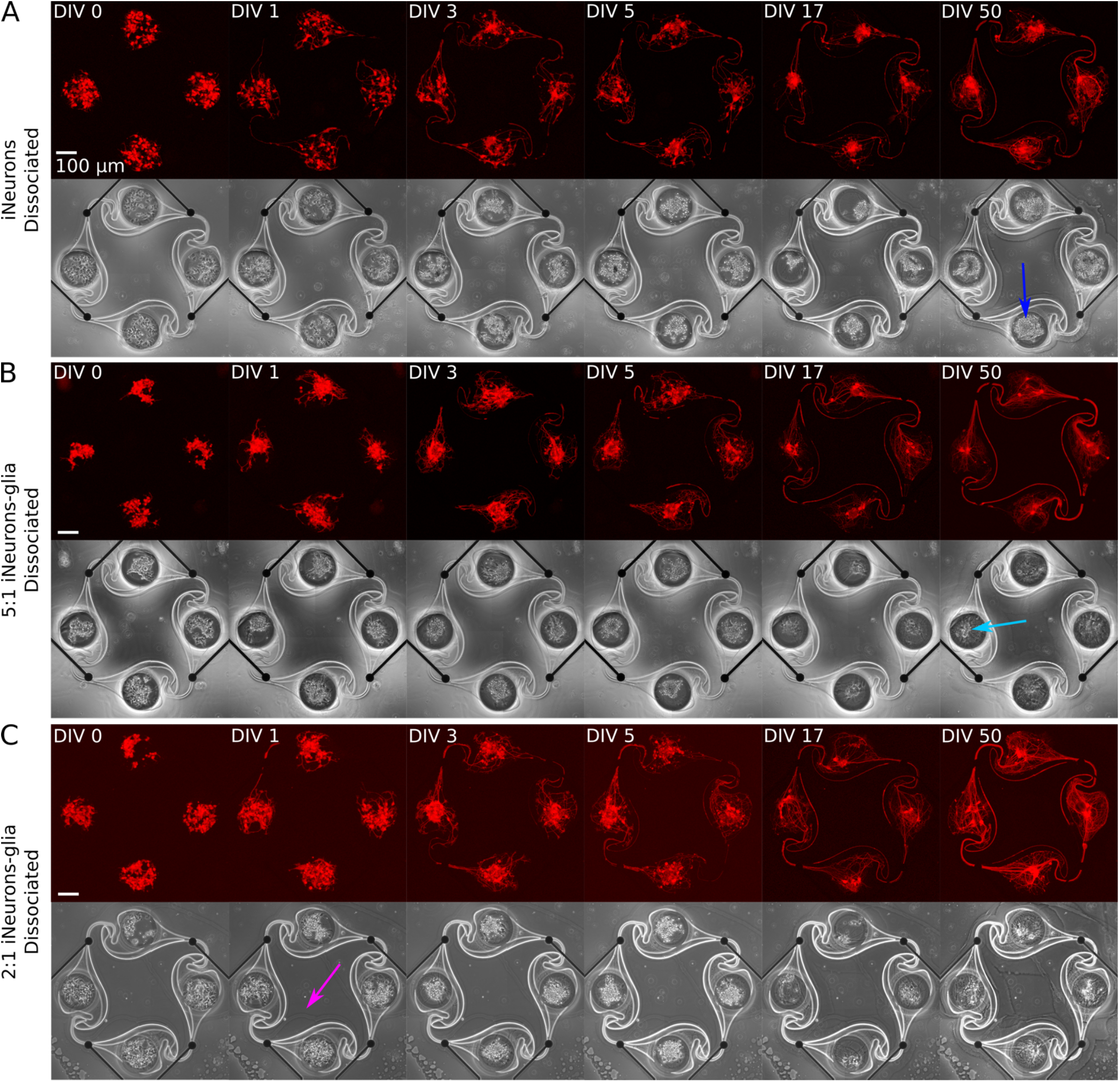
Representative example circuits of iNeurons over time for the 3 different types of dissociated circuits. The red images show RFP-expressing iNeurons and the greyscale image is a phase-contrast view of the circuit, with the edges of the PDMS visible as a bright outline. **(A)** Dissociated iNeurons **(B)** Dissociated 2:1 iNeurons-glia **(C)** Dissociated 5:1 iNeurons-glia. Colored arrows indicate features that are further discussed in the text. ‘DIV’: days *in vitro;* ‘RFP’: red fluorescent protein.

**Figure 6.**
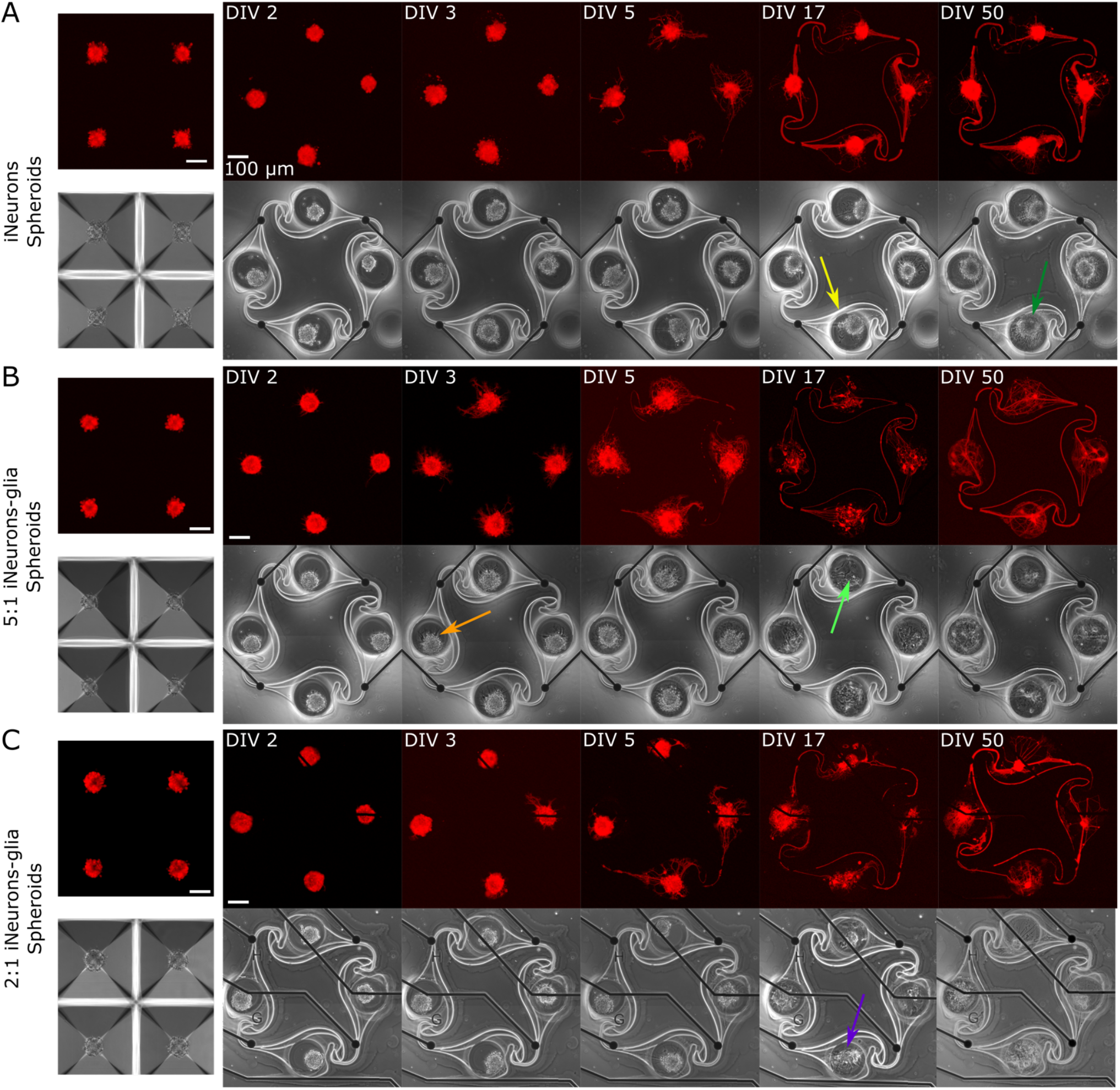
Representative example circuits of iNeurons over time for the 3 different types of spheroid circuits. Spheroids were first formed for two days in anti-adhesive microwells, so example images of spheroids in microwells are shown in A-C. The four spheroids shown on the microwells images are not necessarily the same four spheroids that were seeded into the microstructures on the example images. **(A)** Spheroids of iNeurons. **(B)** Spheroids of 5:1 iNeurons-glia. **(C)** Spheroids of 2:1 iNeurons-glia. Colored arrows indicate features that are further discussed in the text. ‘DIV’: days *in vitro;* ‘RFP’: red fluorescent protein.

Several observations can be made from Fig. 5 and 6. First, the time needed for all four nodes to be connected by axons varies by condition: it takes 3 days for the dissociated iNeurons, 3 to 5 days for the 5:1 dissociated iNeurons-to-glia, more than 5 for the 2:1 dissociated iNeurons-to-glia, and more than 3 days for the spheroid conditions (counting from the day of seeding the spheroids into the PDMS microstructures). At DIV 3, the dissociated glia-containing circuits have a denser network of neurites inside a node, suggesting that at this stage neurites grow inside of the nodes more than along the microchannels. Dissociated iNeurons have less dense neurite growth inside of the nodes, but axons grow faster along the microchannels.

Second, comparing the phase contrast images over time and especially at DIV 50, lines appeared over time along the outline of the PDMS structure and the MEA tracks (an example is highlighted in Fig. 6A with a yellow arrow). This is likely due to the PDMS slightly detaching from the surface. Unlike the lines that appear over time, lines visible in early days phase contrast images *(e.g.* pink arrow, Fig. 5C) are mostly due to a previous use of the MEA. MEAs are typically reused across up to three experiments, but their PDMS structure is replaced with a new one every time. Removing the old PDMS structure might leave a thin residual layer on the MEA’s surface, which is then visible in the phase contrast images.

Third, there is a difference in the amount of dead cells visible between glia and non-glia circuits at later DIVs. On the phase-contrast images of the dissociated iNeurons and iNeuron spheroid circuits, there are many dead cells around and on top of the live cells at all DIVs (e.g. dark blue arrow, Fig. 5A). In the glia-containing conditions, there are also a lot of dead cells in the nodes up to DIV 5, but most of them are gone at DIV 17 and 50 (e.g. light blue arrow, Fig. 5B). The two nodes indicated by the blue arrows can be found in a zoomed-in version in Fig. S5. The removal of dead cells is likely due to the microglia present among the glial cells (see Section 3.1.1). Microglia are immune cells, which can perform phagocytosis to remove necrotic cells. Most of the phagocytosis seems to have taken place between DIV 5 and DIV 17. At DIV 17 and 50, some vacuole-containing, large, flat cells are visible in some of the nodes of the glia-containing circuits (*e.g.* light green arrow, Fig. 6B). This is a typical morphology for activated microglia *in vitro* (Georgieva et al., 2018). Images of the cells growing next to the PDMS microstructures contained cells with a similar morphology (Fig. S4).

Fourth, the cells of the glia-containing spheroid circuits do not present a spheroid morphology anymore at DIV 17 and 50 (e.g. purple arrow, Fig. 5F). iNeuron spheroids are still tightly clustered together, with many dead cells at their surface (*e.g.* dark green arrow, Fig. 5D), whereas the glia-containing spheroids are not visible anymore. The glial cells spread around the node and the remaining iNeurons appear to sit on top of them. Interestingly, at DIV 3 and 5, cells already appear to spread out of the 5:1 iNeuron-glia spheroids (*e.g.* orange arrow, Fig. 5E).

#### 3.3.3 Non-independent circuits were excluded from the electrical activity analysis

Images of the full 15 circuits of each MEA taken at DIV 50 are shown in Fig. S6 and confirm the four trends listed above. Two additional observations can be made: in some circuits, axons inserted themselves under the PDMS; and the glia-containing spheroid circuits had fewer nodes with live iNeurons than the other four conditions. Neurites growing under the PDMS structures often led to a circuit not being compartmentalized anymore, causing it to have potential connections with its neighbor or to the cells located outside of the PDMS microstructures. We termed such circuits “non-independent” (see Fig. 7A and B for examples of independent and non-independent circuits). Images taken at DIV 50 were visually inspected to determine the percentage of independent circuits (Fig. 7C), as well as the percentage of circuits with at least one live iNeuron per node (Fig. S7B).

**Figure 7.**
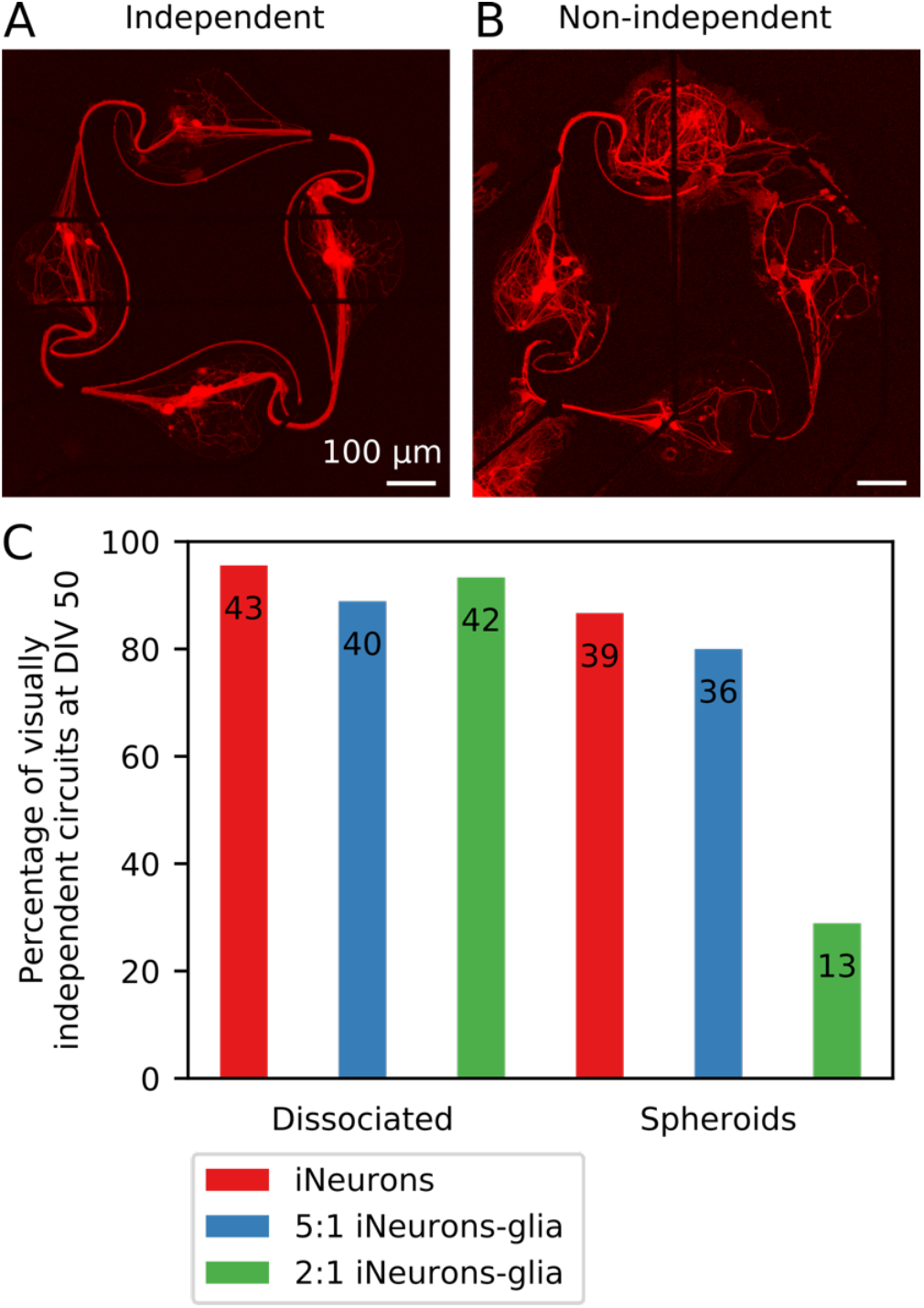
Visually assessed independence of individual circuits for different conditions. **(A)** Example image of a visually independent circuit, where soma and axons stayed in the space intended with the design of the PDMS microchannels (DIV 50). **(B)** Example image of a visually non-independent circuit, where axons grew under the PDMS microstructure and connected the circuit to its neighboring circuits (DIV 50). For both images, the red fluorescence is a genetically encoded RFP expressed by the iNeurons. **(C)** Percentage of independent circuits per condition, assessed visually using individual circuit images taken at DIV 50. The number displayed on each bar corresponds to the number of individual circuits, which are the numbers of circuits used in the analysis displayed in Fig. 8, 10, 11, 12 and 13. For each condition, N = 45 circuits.

In non-independent circuits, axons could insert themselves and grow under the PDMS microstructures due to a compromised adhesion of the microstructure to the MEA. This happened in particular along the tracks of the electrodes of the MEA, as can for example be seen in Fig. 7B. The MEAs used in this work have titanium tracks embedded in a glass carrier substrate and are electrically isolated with a 500 nm layer of silicon nitride deposited by plasma-enhanced chemical vapor deposition (Multi Channel Systems, 2019). Due to the microfabrication process, the tracks are slightly protruding from the surface of the MEA, which is likely the reason why the PDMS microstructures showed poor adhesion around the tracks, leaving space for axons to insert themselves there.

The MEAs seeded with 2:1 iNeuron-glia spheroids mostly had non-independent circuits: only 13 out of 45 circuits (28.9%) did not get connected through axons growing under the PDMS. Two out of three samples had the PDMS microstructures detached over large portions, leading to all but a few of the circuits getting connected together (Fig. S6F). In the 5:1 iNeuron-glia spheroid samples, axons also grew under the PDMS in most circuits (Fig. S6E), but this only resulted in connecting neighboring circuits in 20% of the circuits. In the 3 samples containing spheroids of only iNeurons, there were only a few cases of axons growing under the PDMS, most of which were located at the edge of the PDMS. In addition to the frequent PDMS lifting, glia-containing spheroids had more nodes with no live iNeurons than the other conditions (Fig. S7A). These data suggest that glial cells in spheroids lead to less stable circuits over time.

As we are interested in characterizing the activity of electrically independent circuits, visually non-independent circuits were excluded from the analysis presented in Fig. 8 to 13.

**Figure 8.**
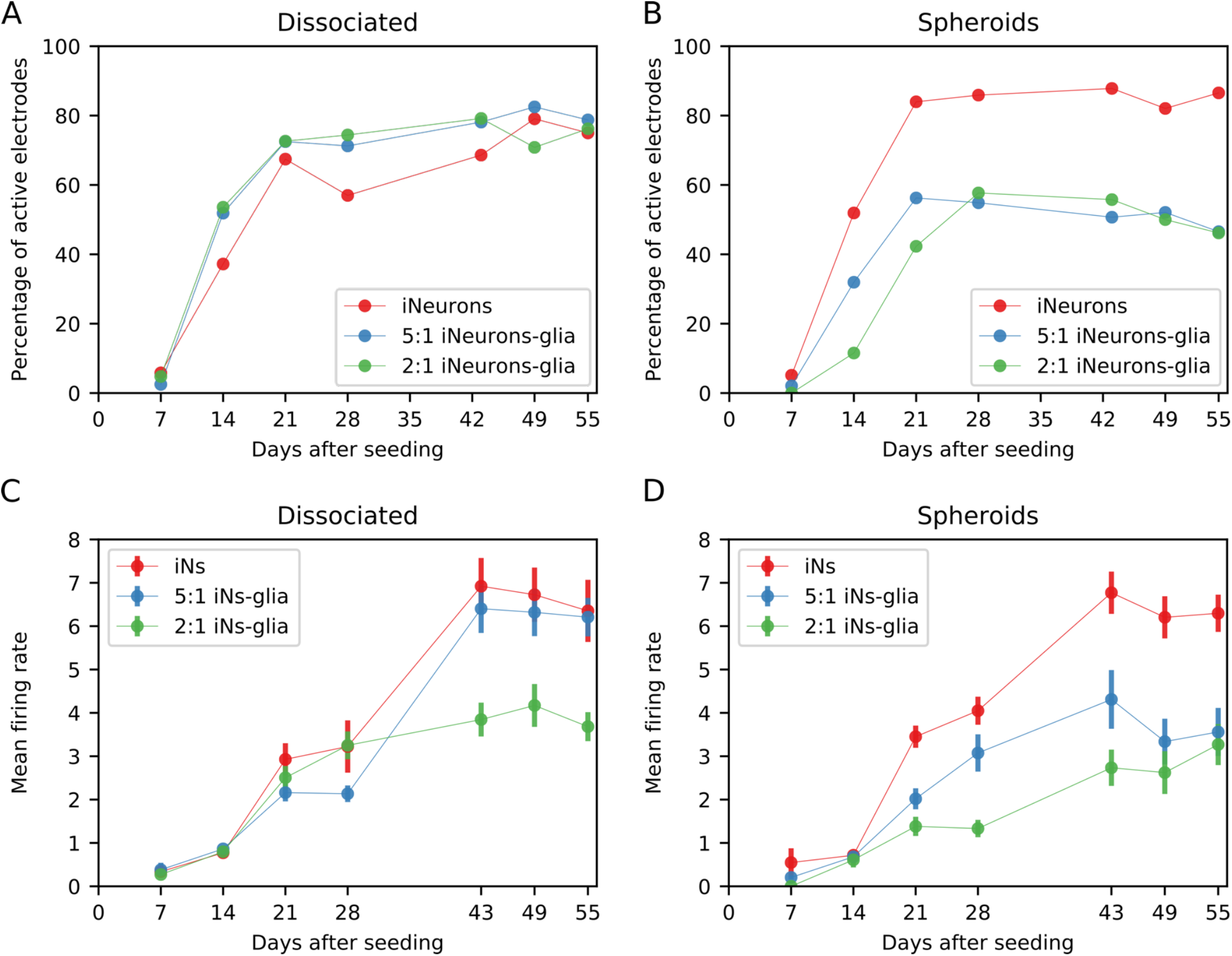
Spontaneous electrical activity for different types of circuits. **(A)** Percentage of active electrodes for different iNeuron-to-glia ratios, for dissociated cells and **(B)** spheroids circuits. Only visually independent circuits were taken into account, so the total number of electrodes is different for each condition and varies between 52 and 172, as shown in Fig. 7C. **(C)** Mean firing rate (MFR) of active electrodes for different iNeuron-to-glia ratios, for dissociated cells and **(D)** spheroids circuits. Error bars represent the standard error of the mean (SEM). ‘iNs’: iNeurons.

**Figure 9.**
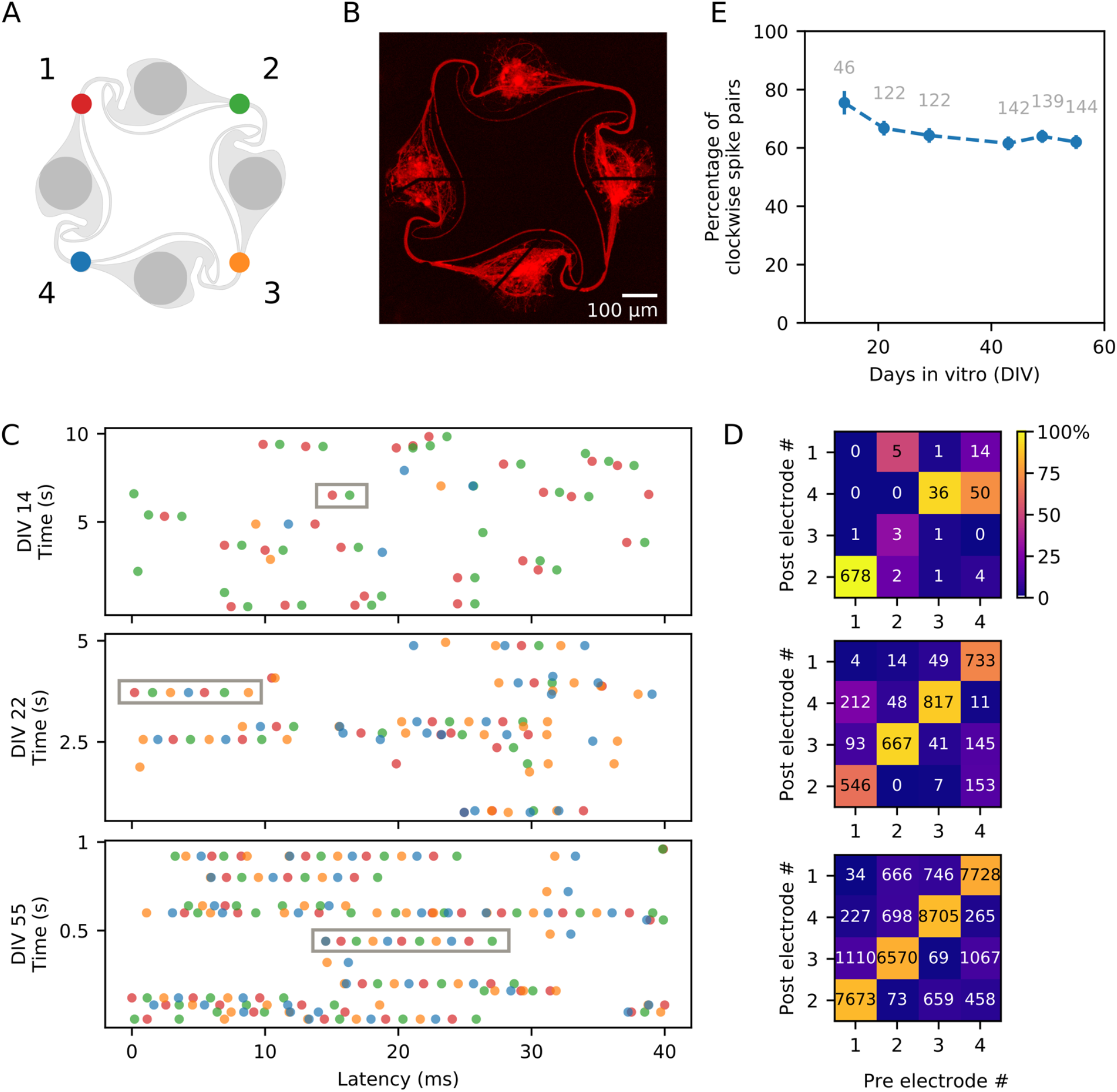
Direction of action potential propagation in the circuits. **(A)** Color code and numbering of the four electrodes of a circuit. Fluorescent image of the circuit of dissociated iNeurons used to generate the example spike trains shown in (C), taken at DIV 50. **(C)** Example spike trains, obtained from the spikes detected on the four electrodes of the circuit shown in (B), with each electrode color-coded as shown in (A). Three different time points are shown: DIV 14 (10 sec long), DIV 22 (5 sec long) and DIV 55 (1 sec long). The spike train (10, 5 or 1 sec long) is split into segments of 40 ms vertically stacked together. The gray boxes show examples of consecutive spikes spaced by less than 5 ms and firing in the clockwise direction (electrodes 1-2-3-4, corresponding to red-green-yellow-blue). **(D)** Frequency map of the occurrence of two consecutive spikes spaced by less than 5 ms, sorted by pre- and post-electrodes. The electrode numbers (1, 2, 3, 4) correspond to the numbering shown in (B). The number displayed on the plot corresponds to the number of spike pairs detected on the pre-/post-electrode pair. The color map corresponds to the percentage of spike pairs from a given pre-electrode that were then detected on a given post-electrode (each column sums to 100%). The spike pairs were detected from 5 min of spontaneous activity recordings on the circuit shown in (A) taken at DIV 14, 22 and 55. **(E)** Percentage of clockwise spike trains, averaged over all the independent circuits of 18 MEA samples. For each circuit, the percentage of “clockwise” spike pairs (electrodes 1 to 2, 2 to 3, 3 to 4, and 4 to 1) was calculated out of the percentage of “clockwise” and “counter-clockwise” spike pairs (electrodes 1 to 4, 4 to 3, 3 to 2 or 2 to 1). Spike pairs occurring between two electrodes with undefined direction (double spike on the same electrode, or spike pair on diagonally opposed electrodes 1 and 3, or 2 and 4) were not taken into account. The numbers displayed in gray correspond to the total number of circuits used for each point. The gray dashed line corresponds to 67.4 %, the expected clockwise percentage of a circuit composed of four nodes with each a 90.6 % probability of clockwise connections. Error bars represent the standard error of the mean (SEM).

**Figure 10.**
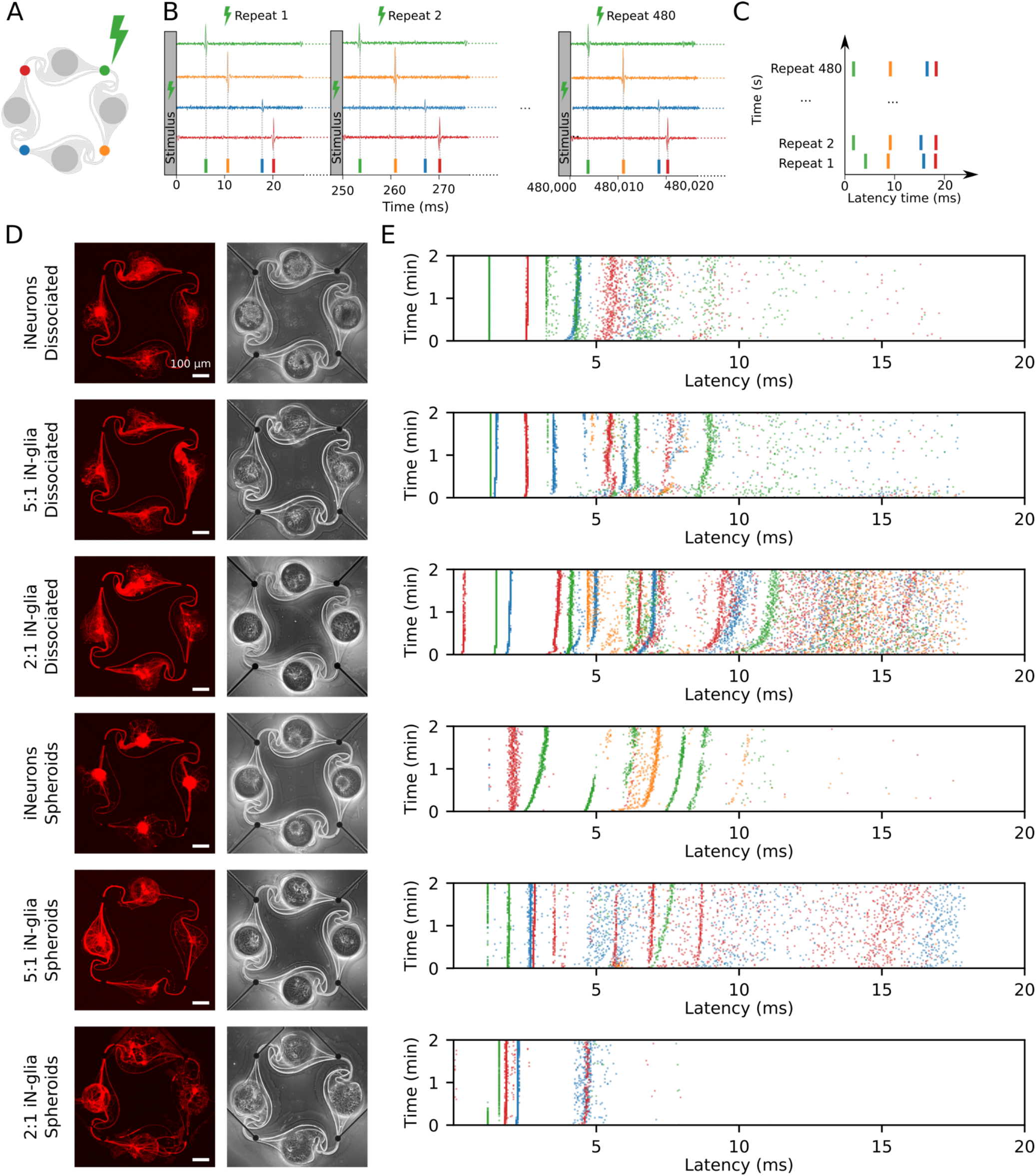
Examples of stimulation-induced electrical activity for the six different types of circuits. **(A)** In the example showed in this figure, an electrical stimulus is applied to the top right (green) electrode of the circuit. The stimulus consists of a 400 μs biphasic square pulse (±500 mV), applied for 2 min at a frequency of 4 Hz (480 repetitions). **(B)** Following the stimulation of the green electrode, spikes are induced and detected on all four electrodes (schematic data, not real data). The detected spikes are color-coded and overlaid as a spike train. **(C)** To represent the data, the 480 repeats of the stimulation-induced spikes are stacked vertically. Spikes that occur with a consistent delay after the stimulus form vertical “bands”. **(D)** Example images of circuits for all six conditions tested, with the RFP-expressing iNeurons (in red) and a phase-contrast image, taken at DIV 50. **(E)** Examples of the stimulation-induced response on all four electrodes of the six circuits shown in (D) upon stimulation of the top right (green) electrode at DIV 49.

**Figure 11.**
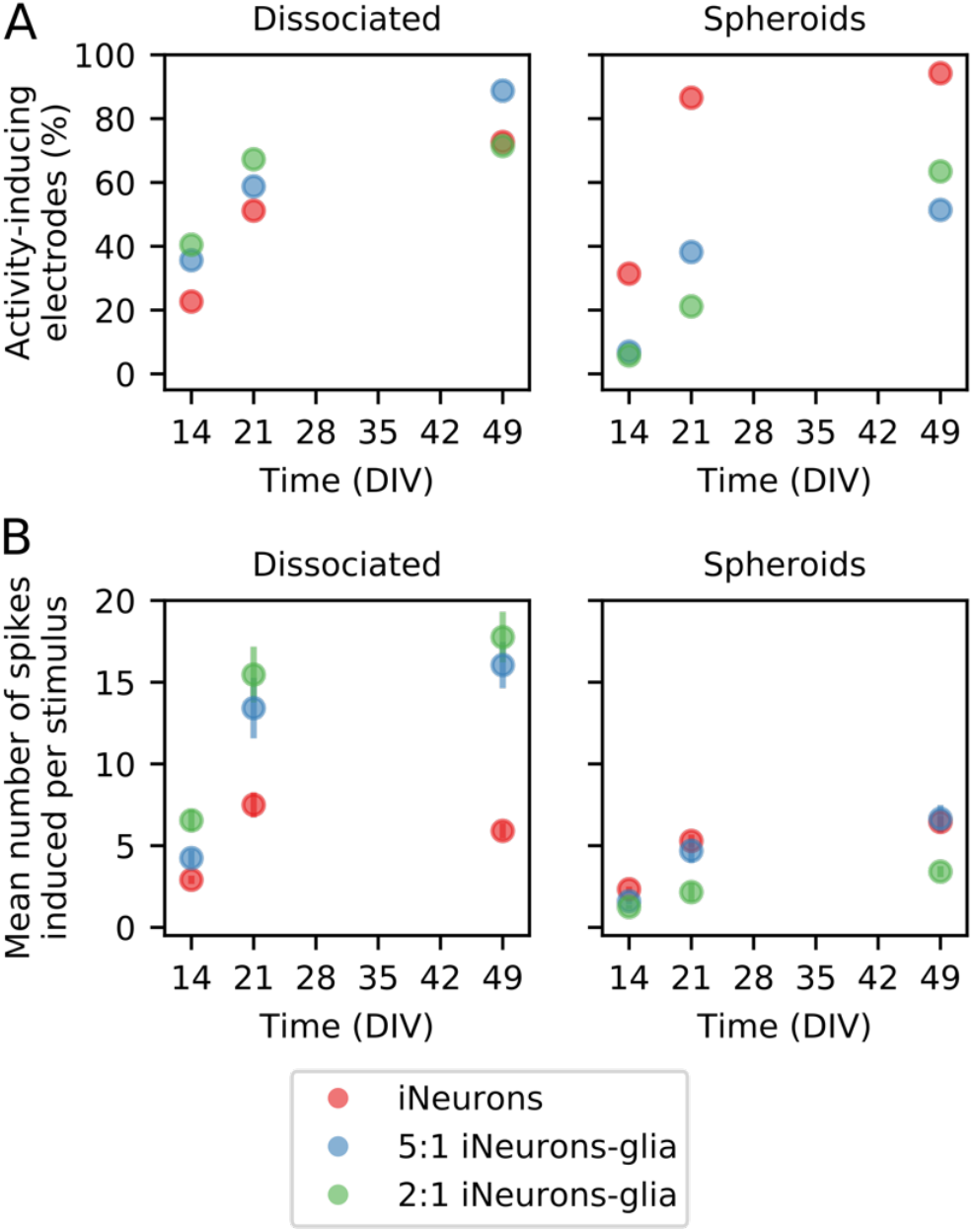
Average response to electrical stimulation across the different types of circuits. **(A)** Percentage of electrodes that induce activity when a stimulus is applied for dissociated (left) and spheroid (right) circuits. An electrode is considered as “activity-inducing”, if at least one spike spike is elicited on average as a response to a stimulus). Only electrodes from independent circuits are taken into account. Therefore, the total number of electrodes vary between N = 52 and N = 176 depending on the condition. **(B)** Quantification of the average number of spikes induced by one stimulus on the “activity-inducing” electrodes plotted in (A). Error bars represent the standard error of the mean (SEM).

**Figure 12.**
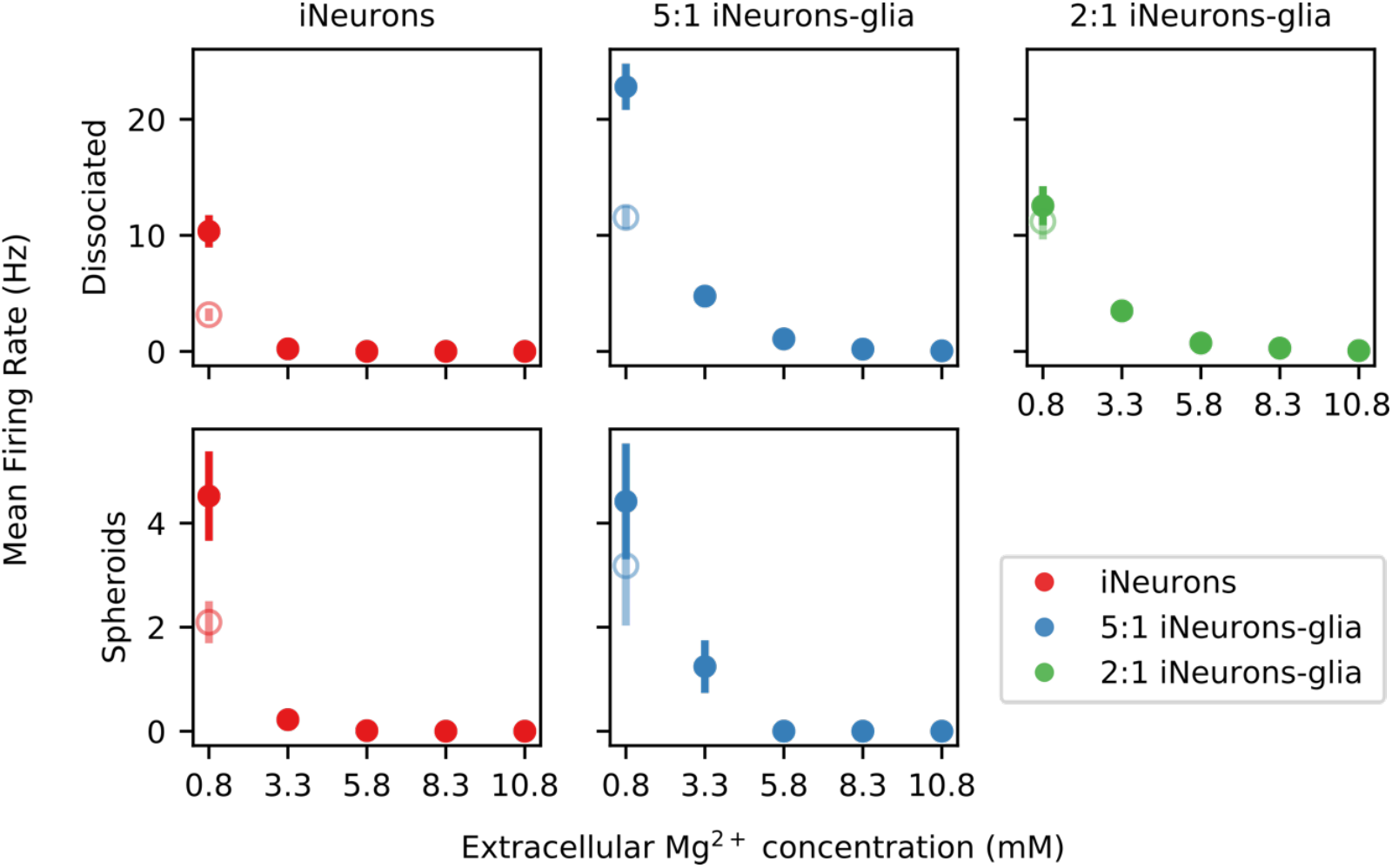
Effect of extracellular Mg^2+^ on the spontaneous firing rate of different types of circuits. Variation of the mean firing rate (MFR) upon sequential addition of magnesium chloride into the culture medium. The empty points represent the recovered post-magnesium MFR, measured after a final medium change. None of the electrodes of the spheroid 2:1 iNeuron-to-glia circuits were active, so no plot is shown for that condition. Error bars represent the standard error of the mean (SEM).

**Figure 13.**
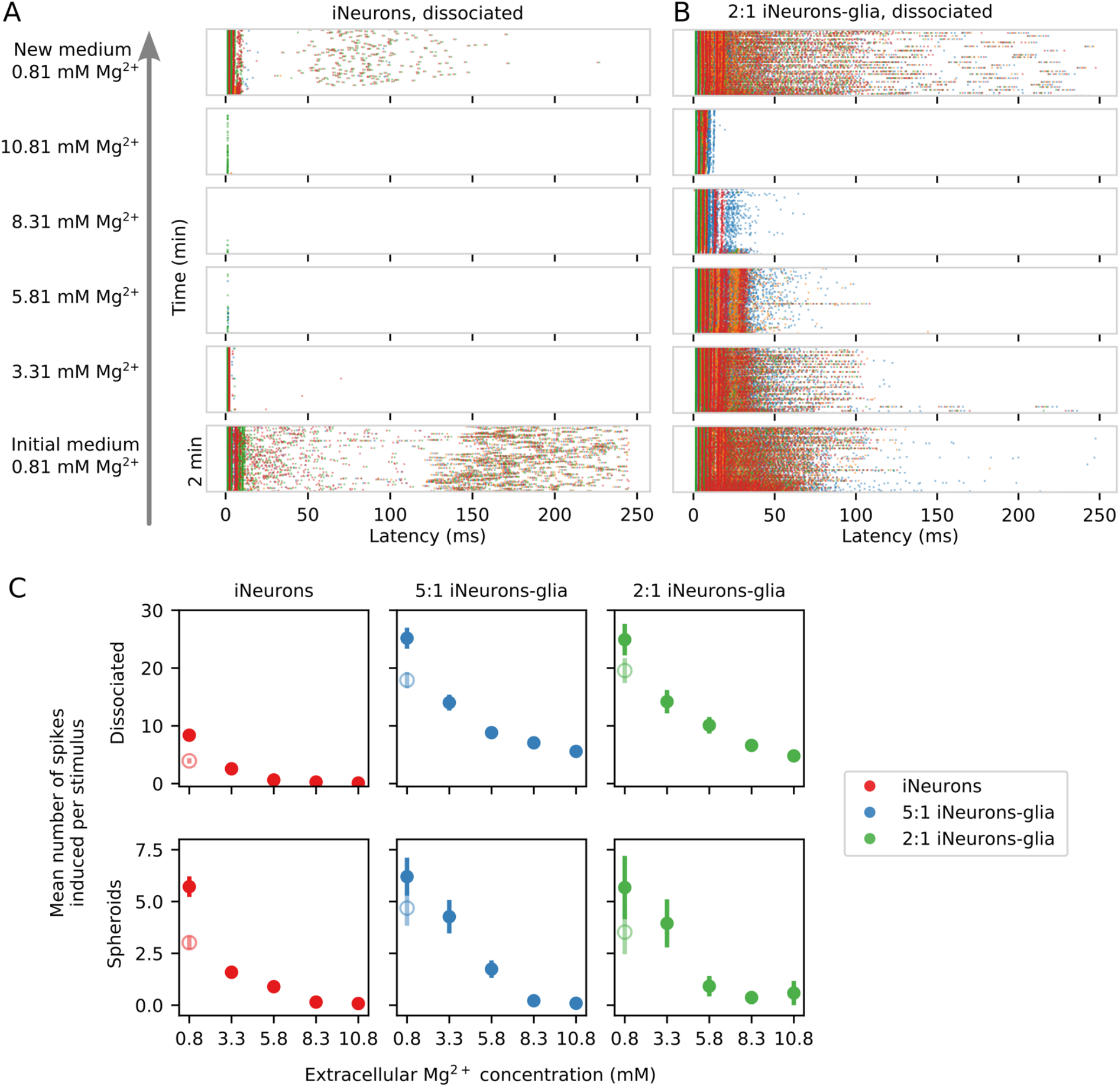
Effect of extracellular Mg^2+^ on the stimulation-induced spiking activity across different types of circuits. **(A)** Example of the changes in the stimulation-induced activity of a circuit of dissociated iNeurons upon sequential addition of magnesium chloride into the culture medium. The stimulated electrode was the top right (green) electrode. **(B)** Example of the changes in the stimulation-induced activity of a circuit of dissociated 2-to-1 iNeurons-to-glia upon Mg^2+^ addition. **(C)** Quantification of the average number of stimulation-induced spikes per electrode upon sequential addition of Mg^2+^ to the medium for the six different types of circuits. The empty points represent the recovered mean number of elicited spikes after the final medium change (post-magnesium addition). Error bars represent the standard error of the mean (SEM).

### 3.4 Circuits of iNeurons and glial cells are electrically active

#### 3.4.1 Active electrodes and mean firing rate of circuits of iNeurons and glial cells

Spontaneous electrical activity was recorded weekly for 5 min on all MEA samples for 8 weeks. The percentage of active electrodes over time was determined for each of the six culturing conditions, taking into account only the visually independent circuits. An electrode was considered active if its firing rate was at least 0.1 Hz. In the three dissociated conditions (Fig. 8A), the number of active electrodes was slightly higher in the two glia-containing conditions up to 42 DIV, after which the number of active electrodes stabilized to slightly under 77% for all three conditions. In the spheroid samples (Fig. 8B), iNeuron spheroids samples consistently had the highest number of active electrodes, with almost twice as many active electrodes than in the two glia-containing spheroid conditions after 55 DIV. This is likely due to the higher number of circuits with empty nodes in these circuits (Fig. S7).

The firing rate of the active electrodes was used to calculate the mean firing rate (MFR) for all conditions. In the dissociated conditions (Fig. 8C), the MFR was similar in all three conditions up to DIV 28. At DIV 43, the MFR of the 2:1 iNeurons-to-glia condition was lower than the 1:0 and 5:1 conditions. In the spheroid conditions (Fig. 8D), the MFR of both glia-containing conditions were lower from DIV 21.

#### 3.4.2 Action potentials predominantly travel in the clockwise direction in the circuits

The design of the PDMS circuit microchannels (Fig. 9A), termed the “stomach” shape, was optimized to guide axons to connect the nodes of a circuit in the clockwise direction (Forro et al., 2018). Based on the observation of hundreds of images of single iNeurons growing in such structures, axons were reported to grow in the clockwise direction between the two nodes of a stomach-shaped circuit in 90.6% of the cases (Girardin et al., 2022). As axons usually transmit action potentials from the axon initial segment to the post-synaptic terminals, controlling the direction of axon growth is expected to influence the main direction of action potential propagation in a network. We thus hypothesize that clockwise axon growth between the four nodes of a circuit should lead to action potentials predominantly propagating clockwise along the four electrodes of a circuit. We verified this hypothesis by analyzing the spike propagation along the electrodes of the circuits.

To analyze the direction of spike propagation in our circuits, we extracted the timestamps of the spikes detected on all four electrodes of each circuit during the weekly spontaneous activity recordings. Only independent and “closed” circuits, with an axon bundle passing on each of the four electrodes (visually assessed), were used in this analysis. Two consecutive spikes were considered to be related, if they were detected within 5 ms of each other on any of the pairs of electrodes in a circuit. Out of four electrodes, there are 16 possible combinations of pairs of “pre-electrode” (where the first spike is detected) and “post-electrode” (where the second spike is detected). The occurrence of two consecutive spikes spaced by less than 5 ms was computed for each combination of pre-/post-electrodes, for all analyzed circuits, over 8 weeks of recordings.

Fig. 9B shows the fluorescent image of one of the circuits from a MEA with dissociated iNeurons. A few seconds of the spike trains detected on this circuit at different DIVs are shown in Fig. 9C, with the spikes color-coded according to the electrode they were detected on. Examples of spike trains for additional DIVs can be found in Fig. S8. The gray boxes highlight sequences of consecutive spikes taking place within 5 ms of each other and firing in the clockwise direction. Over time, the length of such spike sequences tends to increase. Fig. 9D shows a frequency map of consecutive spikes spaced by less than 5 ms for each pair of pre-/post-electrodes within 5 min of recording of the spontaneous activity. The displayed number corresponds to the count of spike pairs on the specific pre-/post-electrode combination, while the color indicates the spike pair frequency relative to the pre-electrode.

Based on the values calculated for each circuit, the overall average percentage of clockwise versus counter-clockwise spike pairs was determined (Fig. 9E). Only the pre-/post-electrode pairs that can be reliably attributed to a “clockwise” or “counter-clockwise” spike propagation were kept to determine the average percentage of clockwise spike trains. Pre-/post-electrode pairs where the direction of the spike propagation is unclear, such as diagonally opposed electrodes (1-3 or 2-4 in Fig.9A), or a repeat of the same electrode, were dropped. The numbers displayed in gray correspond to the number of circuits analyzed at each time point.

The measured percentage of clockwise spike pairs decreases over time, starting at 75.5 % at DIV 14 and dropping to 62.0 % at DIV 55. This decrease is consistent with the observation that axons of iNeurons keep growing over time and do not necessarily stop at their immediate neighbors, complexifying the architecture of the circuits. It is also likely that new synaptic connections form between nodes over time, leading to more complex networks. Overall, the clockwise physical axonal connections resulting from the “stomach” structure is correlated with predominantly clockwise action potential propagation.

#### 3.4.3 Repetitive stimulation of an electrode induces a consistent electrical response on the four electrodes of the circuit

In addition to probing the spontaneous activity of the circuits, we also used electrical stimulation to induce spiking activity. Stimulation was based on a previously reported experimental paradigm (Ihle et al., 2022). The stimulus that we used was a 400 μs biphasic square pulse from 500 mV to −500 mV. This stimulus was sequentially applied for 2 min to each of the four electrodes of a circuit at a frequency of 4 Hz. Therefore, one set of stimuli consisted of 480 repeats of the stimulus on one electrode. Between two sets of stimuli, 30 sec of idle time was left for the circuit to settle. An example of the stimulation-induced activity for one circuit of each of the six conditions is shown in Fig. 10.

Fig. 10A shows the color coding of the four electrodes of a circuit. In the examples shown in this figure, the stimulated electrode was always the top-right, green electrode. Spikes were detected from the data recorded from all four electrodes in the 250 ms time frame following the stimulus onset, for all 480 repeats of the stimulus (Fig. 10B). To visualize the evolution over time of the response to the stimulus, the color-coded stimulation-induced spikes were vertically stacked as shown in Fig. 10C. Spikes occurring at a consistent latency after the stimulus appear as vertical “bands”. Fig. 10D shows the fluorescent and phase contrast images of one example circuit for all 6 conditions tested, imaged at DIV 50. Fig. 10E shows their corresponding stimulation-induced response upon a set of 480 stimuli applied to the top right (green) electrode of each circuit at DIV 49. The morphology of the circuits shown in Fig. 10D is consistent with what was described in Fig. 5 and 6. In the 2:1 spheroid circuit, axons grew under the PDMS. As discussed in Section 3.3.3, this was the case in most of the circuits of that condition. Circuits such as the one shown here, where some axons grow under the PDMS microstructure, but without visible, unwanted connections to a neighboring circuit, were still considered as independent and therefore included in the electrical activity analysis.

When it comes to the stimulation-induced activity, the examples shown in Fig. 10E display vertical bands, *i.e.* a consistent response in the first 10 or up to 20 ms post-stimulus. Similar bands were visible on the stimulation-induced response of all activity-inducing electrodes. In this figure, only the first 20 ms post-stimulus are shown, as it is the time span where the consistent stimulation-induced activity appears to take place. An example of the full 250 ms stacked responses can be found in Fig. S9. As previously reported by Ihle et al. (2022), even in rather simple networks, composed of a few cells and with a controlled topology, the stimulation-induced responses are varied and complex. There is no known relationship between the circuit structure and its stimulation-induced spiking responses.

To compare the different types of circuits, we analyzed the stimulation-induced response of all the electrodes of the independent circuits of each condition (Fig. 11). An electrode was considered as “activity-inducing” if at least one spike was induced on average within the 250 ms time window following the stimulus. Fig. 11A shows the percentage of activity-inducing electrodes per condition. The percentage of activity-inducing electrodes increases over time for all conditions. At all time points, the two glia-containing spheroid conditions have a lower percentage of activity-inducing electrodes than the iNeuron-only spheroids, which is consistent with their low percentage of spontaneously active electrodes (Fig. 8B). For all conditions, the percentage of activity-inducing electrodes at DIV 49 is comparable to the percentage of spontaneously active electrodes at DIV 49 (shown in Fig. 8A and B), but stimulation-induced activity has a later onset (between DIV 14 and 21) than spontaneous activity (between DIV 7 and 14). This is likely due to the fact that spontaneous activity can take place in immature circuits, but stimulation-induced activity is only sustained if functional synapses are present to propagate action potentials between the different neurons of a circuit. Human NGN2 iNeurons were reported to start having functional synapses after around 2 weeks (Zhang et al., 2013).

The mean number of spikes induced per stimulus was calculated for the activity-inducing electrodes and plotted in Fig. 11B. In the dissociated samples, the two glia-containing conditions had a higher mean number of induced spikes. At DIV 49, the mean number of induced spikes is lower in the three spheroid conditions and in the iNeuron dissociated condition than in the dissociated glia-containing conditions. Overall, in the dissociated cultures, the presence of glial cells increases the mean number of induced spikes, but this is not the case for the spheroid cultures.

### 3.5 Measuring the effect of increased extracellular magnesium ions on the electrical activity of the circuits

As a proof-of-concept of the potential of our platform to characterize the effect of an activity-altering compound, we investigated the effect of magnesium chloride on the spontaneous and the stimulation-induced spiking activities of our circuits. Extracellular magnesium ion (Mg^2+^) concentration is known to affect neuronal excitability both *in vivo* and *in vitro*. Due to its high charge density, Mg^2+^ blocks cation channels (Politi and Preston, 2003), which has two main effects. First, it leads to a decrease in the excitability of the membrane and reduces the spontaneous electrical activity of *in vitro* neuronal cultures (Crain et al., 1968). Second, it impairs the release of excitatory neurotransmitters, in particular by blocking voltage-gated calcium channels (Cuciureanu and Vink, 2011). Under physiological conditions, Mg^2+^ is also a specific antagonist of N-methyl-D-aspartate (NMDA) receptors (Nowak et al., 1984).

In the Neurobasal-based medium used in the presented work, the baseline concentration of Mg^2+^ is 0.81 mM. We sequentially increased the concentration of Mg^2+^ in the medium up to 10.81 mM, by steps of 2.5 mM, using a concentrated solution of magnesium chloride. This was done for one MEA for each of our six culturing conditions. The spontaneous electrical activity and the stimulation-induced activity were recorded in the initial medium, after each Mg^2+^ addition step, as well as after a final complete medium exchange back to the original culture medium. These recordings were performed between DIV 56 and 58.

#### 3.5.1 Spontaneous electrical activity upon magnesium addition

Fig. 12 shows the MFR upon addition of extracellular Mg^2+^ for five of the six types of tested conditions. On the 2:1 iNeurons-to-glia spheroid MEA where extracellular Mg^2+^ was added, none of the independent circuits had active electrodes, which is why it is not represented on the figure. The MFR after the final medium change is shown as an empty circle and is lower than the initial MFR in all conditions.

Consistent with what is reported in the literature, increasing the extracellular Mg^2+^ decreases the spontaneous activity of the networks. In both the dissociated and the spheroid cases, the samples with only iNeurons were more sensitive to Mg^2+^ than the glia-containing samples, with a lower MFR upon a 3.3 mM extracellular Mg^2+^ concentration. It is possible that the glial cells, in particular the astrocytes, play a buffering or a protective role in the presence of elevated Mg^2+^ concentrations.

#### 3.5.2 Stimulation-induced activity

The stimulation-induced activity is also affected by the increase in extracellular Mg^2+^. Two representative examples of the stimulation-induced response over 250 ms upon sequential addition of extracellular Mg^2+^ can be seen in Fig. 13A (dissociated iNeurons) and B (dissociated 2:1 iNeurons-to-glia). The glia-containing sample had a higher number of induced spikes at all concentrations of Mg^2+^. This was verified by quantifying the average number of stimulation-induced spikes over all the independent, activity-inducing electrodes of each of the 6 MEAs. Results can be seen in Fig. 13C, with the empty circles representing the mean number of induced spikes after the final medium change. The post-magnesium recovered activity was lower than the initial activity for all conditions. Generally, the response to increased extracellular Mg^2+^ concentrations followed a similar trend in the two iNeurons samples (dissociated and spheroid), in the two glia-containing dissociated samples (5:1 and 2:1), and in the two glia-containing spheroid samples (5:1 and 2:1). Measurements were performed over 3 days, recording from two samples per day.

In the two iNeuron samples and in the two glia-containing spheroid samples, the induced activity was almost fully suppressed with an extracellular Mg^2+^ concentration of 8.3 mM. In general, stimulating the neurons rather than merely recording their spontaneous activity shifted the threshold of extracellular Mg^2+^ necessary to shut down all electrical activity in the circuits. This is to be expected, since stimulation should elicit action potentials in the close proximity of the stimulated electrode at a much higher rate than when relying on spontaneous activity. Stimulation seems to partially overcome the magnesium-induced decrease in spontaneous activity. Consistent with what was observed in the spontaneous activity recordings (Fig. 11), the presence of glial cells in the two spheroid samples made the cultures less sensitive to the presence of extracellular Mg^2+^ upon stimulation-induced activity. Overall, the results presented in Fig. 12 and 13 both support the hypothesis that the presence of glial cells decreases the sensitivity of the cultures to extracellular Mg^2+^.

In the two dissociated glia-containing samples, spikes were still induced in the first few ms after stimulation even at the highest concentration of extracellular Mg^2+^ (10.81 mM), as illustrated in Fig. 13B and in the zoom-in of the first 20 ms of response after stimulus shown in Fig. S10. This suggests that in these samples, the presence of glial cells allows for sustained activity even at high Mg^2+^ concentrations. The spikes suppressed by the presence of increased Mg^2+^ are the ‘late’ spikes, taking place later than 15 ms after the stimulus and likely mediated by synaptic connections between the neurons of a circuit. Synaptic transmission is blocked in the presence of high Mg^2+^. The ‘early’ spikes visible in the first 15 ms post-stimulus might either be a direct consequence of the stimulus, or rely on synaptic connections that are not fully inhibited by the presence of Mg^2+^. The dissociated glial-containing samples respond differently than the spheroid glial-containing samples to the presence of magnesium. The level of stimulation-induced activity was lower in the spheroid samples even without added Mg^2+^ and the samples usually had more circuits with empty nodes, which might explain the observed differences.

#### 3.5.3 Comparing the spontaneous and stimulation-induced activities

The MFR is known to be sensitive to several environmental factors, such as a temperature or humidity change, moving the cultures, or differences in ionic concentrations. A culture left in the incubator for a few days will undergo medium evaporation, leading to a slightly increased ionic concentration. In general, the vulnerability of the MFR to environmental factors is a limitation of that measure. To inspect the stability of the MFR compared to the stimulation-induced activity, we sequentially measured the spontaneous and stimulation-induced activity in two samples after four medium changes and after addition of a small volume (5 μL) of ultrapure water and DMSO. Results are shown in Fig. S11. The spontaneous MFR was very sensitive to medium changes, with a coefficient of variation of around 45 %, whereas the stimulation-induced activity was more stable, with a coefficient of variation of about 11 %. Overall, the stimulation-induced response is less dependent on environmental factors and more stable. It is therefore a more reliable metric than the spontaneous MFR.

## 4 DISCUSSION

The results presented in this work support three main findings: first, the PDMS microstructure-based platform is very modular; second, the presence of glial cells impacts the morphology and the electrical activity of the iNeuron circuits; and third, the platform has potential for drug testing.

### 4.1 A highly modular platform

The PDMS microstructure-based platform was successfully used to build topologically controlled circuits of human-derived iNeurons and rat primary glial cells at different ratios, as either dissociated cells or spheroids. The system is compatible with imaging assays, as well as with MEA recording and stimulation, including the addition of activity-altering compounds (here magnesium ions) into the cell medium. Three elements from our results are of particular interest when it comes to the modularity of the platform: the stable antifouling coating, the control over the direction of the information flow, and the possibility to use spheroids in the platform.

#### 4.1.1 PDMS antifouling coating

We showed that coating the PDMS microstructure with PFPA-PVP leads to a very efficient and resistant antifouling coating (Fig. 4). This makes the system more robust when using neurons or cells that tend to adhere to uncoated PDMS. In addition, because the adhesion of PFPA-PVP relies on UV light exposure, instead of a uniform antifouling coating, it would be possible to pattern specific antifouling areas on top of the PDMS using a UV mask (Weydert et al., 2019). Non-patterned areas could either be left uncoated, or be coated with a neuron-adhesive molecule such as PDL or laminin. This would add a layer of modularity to the platform, enabling additional connections between the circuits and allowing to build circuits with connections that could otherwise not be engineered using only a 2D plane.

#### 4.1.2 Direction of information flow

We demonstrated that physically constraining axons to grow in a preferred direction led to action potentials mainly propagating in the corresponding direction (Fig. 9). This allows building circuits with less variations and with a more ordered topology, which could for example be important when trying to recapitulate realistic *in vivo* brain circuits. Networks with a simpler structure should also be easier to understand and to model *in silico*. Much effort has been put into linking the structure and functionality of random *in vitro* networks of neurons on MEAs (Ullo et al., 2014; Poli et al., 2015; Pan et al., 2015). Because of the low number of cells in our system, the structure of the networks can easily be determined by fluorescent microscopy. In the present work, RFP-expressing iNeurons were used, allowing to visualize the neurons’ whole morphology. To get more information about the network’s connections, additional genetically encoded fluorescent markers could be added, for example to locate synapses using the presynaptic terminal marker PreSynTagMA and the postsynaptic marker PSD-95 (Mateus et al., 2022). This should make it possible to precisely link the structure and electrical functional activity of the network.

#### 4.1.3 Recording electrical activity of spheroids

Our system can reliably direct the bundles of axons growing between spheroids placed in the different nodes of a circuit, enabling to record the information that they exchange and to stimulate them effectively. This is an interesting possibility, because in the absence of PDMS microstructures, recording from a spheroid is usually limited to the cells located at the bottom of the spheroid. Using PDMS microstructures to force the axons of a spheroid to be in close proximity to the recording electrodes might open the way to getting a better insight into their electrical activity.

#### 4.1.4 Possible improvements for the platform

##### 4.1.4.1 A better protocol for spheroids of iNeurons and glia

While spheroids of iNeurons could be used reliably to build networks, results obtained with glia-containing spheroids were limited. The two reasons for that are that the PDMS microstructure detachment was observable in most circuits of glia-containing spheroids, and that these circuits had more nodes where no cells survived than the other conditions. PDMS detachment could be prevented by plasma bonding the PDMS to the MEA surface or mitigated by using MEAs with a flatter surface to avoid the decreased adhesion of PDMS along the MEA tracks.

However, the poor health of the spheroids points towards other potential problems, possibly an inappropriate spheroid generation protocol or the lack of space for the high number of cells seeded into the structures. The spheroid generation protocol was based on Cvetkovic et al. (2018), replacing iPSC-derived astrocytes with rat primary glial cells. Primary cells might not be a suitable cell type to form spheroids. Therefore, replacing them with iPSC-derived astrocytes could improve the efficiency of the protocol. Better nutrient diffusion could be addressed by redesigning the PDMS microstructure to have larger node diameters (currently 170 μm) and wider microchannels (currently 10 μm) to allow for proper nutrient diffusion. The height of 4 μm of the microchannels should be kept unchanged, as it prevents soma from migrating into the channels.

##### 4.1.4.2 Adapting the PDMS microstructure design

A particularity of the iNeurons is their capacity to grow several mm over the course of an experiment. The “stomach” PDMS structures were originally designed for use with cortical and hippocampal rat primary neurons (Forro et al., 2018), which typically grow by 1-1.4 mm over 4 weeks *in vitro* (Kaneko and Sankai, 2014). By contrast, NGN2 iNeurons grow by 3.7 mm on average in the same time interval (Rhee et al., 2019). The shortest path between the centers of two nodes of a circuit is about 680 μm, which corresponds well to the typical axonal length of a rat cortical or hippocampal neuron, but is much shorter than the average length of an iNeuron axon. The tendency of iNeuron axons to grow to the top of the PDMS and to form networks of axons there, is an indication that they need more space. For that reason, the aforementioned redesign of the PDMS structure could also include longer microchannels.

### 4.2 Effect of glial cells on iNeurons circuits

#### 4.2.1 Phagocytosis of dead cells

The presence of glial cells has an important impact on the morphology of the circuits: in the glia-containing circuits, dead cells got phagocytosed after about 2 to 3 weeks (Fig. 5), very likely due to the presence of microglia (Fig. 2). While protocols exist to obtain pure cultures of astrocytes (Uliasz et al., 2012), in the presented work, we were interested in using a mixture of glial cells in the circuits. To explore the role of specific subtypes of glial cells on the iNeurons circuits in more detail, it would be possible to use purified populations of either primary astrocytes or primary microglia. We did not verify nor discuss the presence of oligodendrocytes, a third type of glial cells responsible for the myelination of axons *in vivo*.

#### 4.2.2 Spontaneous electrical activity

Astrocytes are known to perform glutamate clearance at synapses to prevent neuronal hyperexcitation (Mahmoud et al., 2019). They might play a similar role *in vitro.* The MFR of iNeuron cultures is rather high (Fig. 9, around 6-7 Hz at DIV 43), but seemed to be lower in the presence of glial cells (2-4 Hz) for three out of the four types of iNeurons/glia circuits.

#### 4.2.3 Functional synapses

Several observations support the presence of functional synapses in the circuits. First, after three weeks in culture, spontaneously elicited action potentials often propagate over the electrodes of a circuit in sequences lasting for more than 10 ms (Fig. 9). Second, in many circuits, stimulating one of the electrodes of a circuit induces sustained and complex activity over more than 10 ms (Fig. 10 and Fig. S9). Third, the addition of magnesium mostly blocks sustained activity (Fig. 13). These effects were observed in all samples regardless of the presence of glial cells, which means that synapses can form even in the absence of astrocytes. Astrocytes were reported to accelerate the onset and number of synapses when co-cultured with iPSC-derived neurons (Johnson et al., 2007). In our circuits, while the presence of glial cells did not have a visibly high impact on the onset of spontaneous activity (Fig. 8C and D), it increased the stimulation-induced activity in dissociated circuits from 3 weeks in culture (Fig. 11). This suggests that the presence of glial cells facilitates synapse formation in dissociated cultures. Finally, the presence of glial cells lowered the sensitivity of iNeurons to high extracellular Mg^2+^, suggesting that they may provide a buffering or a protective effect (Fig. 12 and 13).

### 4.3 A platform with drug screening potential

This work provides a proof-of-concept that PDMS microstructure-constrained small circuits of neurons on MEAs can be used to test the effect of soluble compounds on the spiking activity of the circuits, which could in turn be used for drug screening applications. The compatibility of the platform with imaging assays such as live-dead staining would also allow to test for the toxicity of a molecule added to the culture. Here we discuss how the PDMS microstructure-constrained circuits compare to random networks, the potential of spheroids for drug screening applications, and the use of stimulation-induced activity as a readout.

#### 4.3.1 PDMS-constrained networks vs random networks

*In vitro* random networks of neurons have been used in combination with MEAs for drug screening applications, in particular using multi-well MEAs (Kim et al., 2014). We propose to use PDMS microstructures to constrain neurons into small networks, as this approach presents several advantages compared to random networks of neurons. First, there is no cell detachment over time and circuits are stable over months. This gives the possibility to use the platform with cell types that need time to mature, *e.g.* to recapitulate disease-specific phenotypes (Odawara et al., 2016). Second, the microchannels improve the electrical recordings, which have high signal-to-noise ratio (FitzGerald et al., 2008; Pan et al., 2014) and can be linked to the group of neurons from which they arose. The electrical stimulation is also specifically targeted to the bundle of axons located on top of the electrodes, giving the possibility to measure specific stimulation-evoked responses (Fig. 10). Third, medium can be changed with relatively little disturbance to cells compared to random cultures, as they are protected from turbulent flow by the thickness of the PDMS microstructures. With a diameter of 170 μm and a height of 120 μm, each node contains roughly 2.7 nL. The cell medium volume used in a typical MEA is 1 mL. Upon complete medium change, the entire medium is aspirated from the MEA, except for a thin film of liquid of a few μL. In comparison, the volume left in the nodes is negligible. Overall, we estimate that a careful medium change should allow for a rapid dilution of at least 1:1000 of the compound added to the medium, while leaving the circuits mostly undisturbed. Fourth, unlike in random cultures, a very small number of cells is needed to build circuits, making it compatible with the use of rare or expensive cell types, as long as survival at low cell density is possible. Each sample is composed of 15 circuits produced under similar conditions, which allows to compare parallel repeats of the circuits. The main disadvantage of the PDMS microstructure platform compared to a multi-well MEA is its lower throughput. Given the advantages presented with regards to random cultures, but considering the lower throughput in comparison with multi-well MEAs, the platform could for example be used in combination with drugs that were found to have an effect on neural cultures on multi-well MEAs, in order to study the compounds in more detail.

#### 4.3.2 Spheroids

Given the interest in 3D systems for drug testing, we hypothesized that the use of spheroids in our platform would be a better approach than the use of dissociated cells (Jensen and Teng 2020). We expected functional differences between dissociated and spheroid conditions, as spheroids should allow for cell-cell interactions that are normally more constrained in 2D cultures (Cvetkovic et al., 2018). However, the conclusiveness of the results obtained with spheroids is limited. As mentioned in Section 4.1.4, the iNeuron/glia spheroid protocol needs to be improved, as the spheroids did not retain their shape and showed poor viability. In contrast, the iNeuron spheroids kept their spheroidal shape throughout the 8 weeks of culturing. However, they did not present very different electrical characteristics from dissociated iNeurons. Our results indicate that more work is needed to find a proper culture protocol for iNeuron/glia spheroids in PDMS microstructures and that dissociated cells might be a sufficient model for the use of the system in early phases of drug screening.

#### 4.3.3 Stimulation-induced activity as a readout

Using stimulation to induce activity at a controlled rate is a more reliable readout system than relying on spontaneous action potentials. The presence of spontaneously recurring spike sequences (Fig. 9C) suggests defined connection paths between the neurons in our circular networks. Using stimulation to induce activity increases the occurrence of such recurring spike sequences (Fig. 10E and Fig. S9). In addition, a compound can be added to the medium at a precise concentration to measure its effect on the stimulation-induced responses (Fig. 13). Finally, our data suggests that stimulation-induced activity is more stable and consistent over time and contains more information compared to spontaneous activity (Section 3.5.3).

## 5 CONCLUSIONS AND OUTLOOK

In this work, we built upon the existing reports of a versatile and modular PDMS-based platform for bottom-up neuroscience research (Forro et al., 2018; Ihle et al., 2022; Girardin et al., 2022; Duru et al., 2022). We further demonstrate the versatility of the platform to build circuits of different ratios of human iNeurons and rat primary glial cells, initially seeded as either dissociated cells or spheroids, which can be cultured, imaged, and electrically monitored for more than 50 DIV. As a proof-of-concept for the use of such a platform for pharmacological molecule screening, we evaluated the effect of the sequential addition of magnesium chloride on the spontaneous and stimulation-induced electrical activity of the different types of circuits. Because it is compatible with iPSC-derived neurons, the platform could be used with patient-derived cells or with neurons genetically engineered to recapitulate a disease phenotype (Xu and Zhong, 2013). Instead of primary glial cells, astrocytes of human origin, such as iPSC-derived astrocytes or primary human astrocytes, could be used.

While the results obtained are promising, several developments are still needed to make this platform compatible with early drug discovery. In particular, the platform should allow for higher throughput, be interfaced with medium-exchange pumps, and a set of data analysis tools should be developed to further analyze the stimulation-induced activity. The current throughput is limited by the 15 circuits per MEA. Therefore, MEAs with a higher number of electrodes should be considered. Adding a perfusion system to the MEA would allow continuous and controlled addition of molecules to the medium, which should minimize the effects of external factors such as MEA transport and temperature changes on the stimulation-induced activity. In addition, a perfusion system would enable automatized medium change and uninterrupted long-term recordings. Finally, the collected stimulation-induced data is rich in information, but the tools to analyze such data are lacking. The magnesium ions used in the present work have comparatively large and non-specific effects on the networks’ electrical activity, and are thus easy to detect. An automated band detection algorithm would increase the sensitivity of our model system to lower concentrations and thereby also to more specific effector molecules that only target a certain subtype of ion channels.

When all of these aspects are integrated into the platform, it should be possible to elicit and subsequently classify characteristic concentration-dependent effects of molecules based on their known mode-of-action (e.g. NMDA receptors antagonists and AMPA receptors antagonists). New molecules could then be classified based on their effect on the networks response behavior and thereby predict the molecular targets. Overall, the system presented here has the potential to become a drug-testing platform to find new molecules of interest for the treatment of neurological diseases.

## AUTHOR CONTRIBUTIONS

SG and JV designed the research project, with inputs from SJI, JD and TR. SG wrote the manuscript with support from all co-authors. AM, MK, and LT performed preliminary experiments. SG conducted the experiments with support from MK. SG analyzed the data, with the help of SJI for the electrophysiology data. JV secured funding for the projects. IF and MM produced the iPSCs and iPSC-derived neurons. All co-authors reviewed and approved the manuscript.

## ACKNOWLEDGMENTS

Thank you to Sean Weaver for advice regarding cell culture and microscopy, as well as to Sinéad Connolly for proof-reading the manuscript.

