## Supplementary Material for "Engineering circuits of human iPSC-derived neurons and rat primary glia"

#### 1 IMMUNOFLUORESCENT STAINING OF ASTROCYTE CULTURE: MAP2

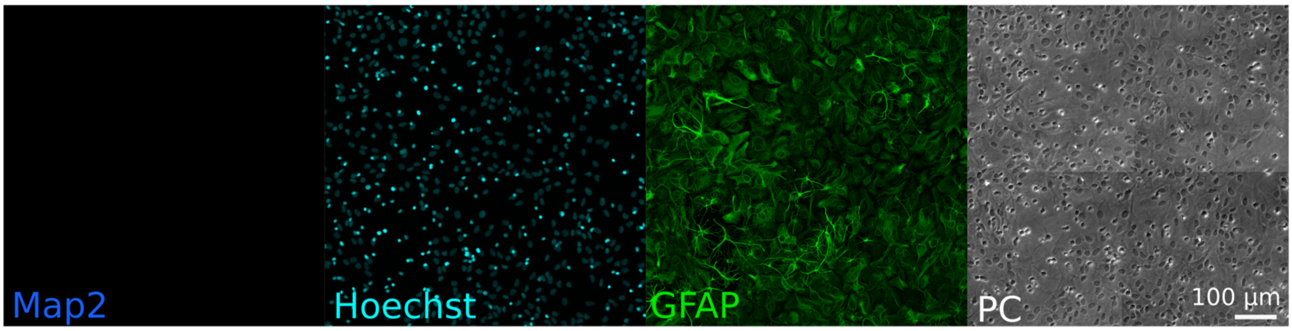


**Figure S1. Immunofluorescence staining of rat primary glial cells cultures.** MAP2 (blue) stains neurons, Hoechst (cyan) stains nuclei, GFAP (green) stains astrocytes. 'PC': phase contrast (brightfield) images. All channels but MAP2 are the same as shown in Fig. 2A. No MAP2 staining is visible, confirming the absence of rat primary neurons in the cultures of rat primary glial cells.

#### 2 LIVE-DEAD/HOECHST STAINING

##### 2.1 Materials & Methods: Live-dead/Hoechst staining

A solution of 2 µM of Calcein-AM and 8 µM of ethidium homodimer-1 (both from L3224, Thermo Fisher) in DPBS (14190-144, Thermo Fisher) was incubated with the sample for 12 min. The same volume of a solution of 2 µM of Hoechst 33342 (H3570, Thermo Fisher) was added to the sample and incubated for another 8 min. The sample was then carefully washed once with DPBS and left in warm DPBS for imaging.

##### 2.2 Results: Live-dead and Hoechst staining

Based on the nuclei stains, it is clear that the cell bodies are located in the nodes of the circuits, while green-labelled axons grow through the microchannels. The dissociated circuit of iNeurons (A) appears to have a less dense axon network than the glia-containing circuit (B), but based on the number of live nuclei visible (bright blue spots), it also contains fewer live cells. In the spheroid circuits, the nuclei stain shows that iNeurons stay tightly clustered even after 24 DIV (C), whereas the glia-containing spheroids spread around more (D). Phase contrast images revealed that in glia-containing spheroid samples, it is mainly the glial cells that spread around to form a layer at the bottom of the node, on which lie the iNeurons, still clustered together. On the live stain spheroid circuit images, it is also visible that axons inserted themselves under the PDMS, for example on the right hand side node of both the iNeuron and the 5:1 iNeuron-to-glia circuits. Such PDMS detachment was not usually observed in the dissociated samples and is probably due to the high number of axons present in the spheroid samples. To avoid unwanted PDMS lift by axons, the number of cells per spheroid was reduced by half in the electrophysiology experiments presented in the next figures. Finally, with the phase-contrast and red stains, a rather high number of dead cells are visible.

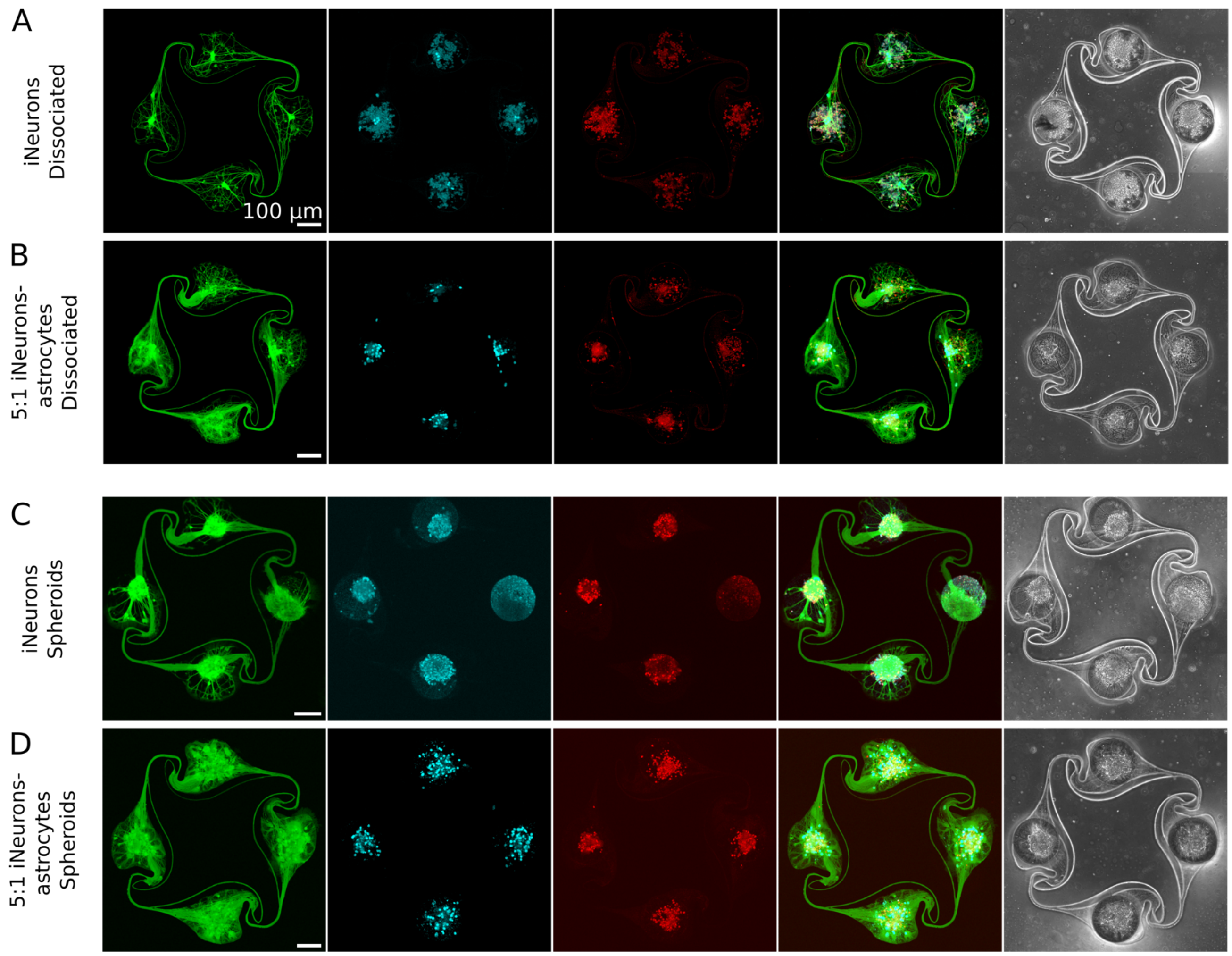


**Figure S2.** Representative examples of circuits of **(A)** dissociated iNeurons, **(B)** 5-to-1 iNeurons-astrocytes (dissociated), **(C)** spheroids of iNeurons and **(D)** spheroids of 5-to-1 iNeurons-astrocytes. For all conditions, both iNeurons and astrocytes are labeled with the stains. The first column (green stain, Calcein AM) shows live cells. The second column (blue stain, Hoechst) shows the cell nuclei. The third column (red stain, ethidium homodimer-1) shows dead cells. The fourth column shows a merge of the first three columns. The fifth column shows a brightfield, phase contrast picture of the circuit.

#### 3 UNWANTED AXON GROWTH ON PFPA-PVP COATING

PDMS microstructures coated with the PFPA-PVP coating could effectively prevent unwanted axonal growth on top of the PDMS for more than a month (Fig. 4). Out of the 12 repeats of PFPA-PVP-coated PDMS microstructures, one of them showed extensive unwanted axonal growth. Images of the top surface of that microstructure are shown in Fig. S3. The six circuits showed on the image presented unwanted axonal growth. The network of axons connects with the cells located outside of the PDMS microstructure.

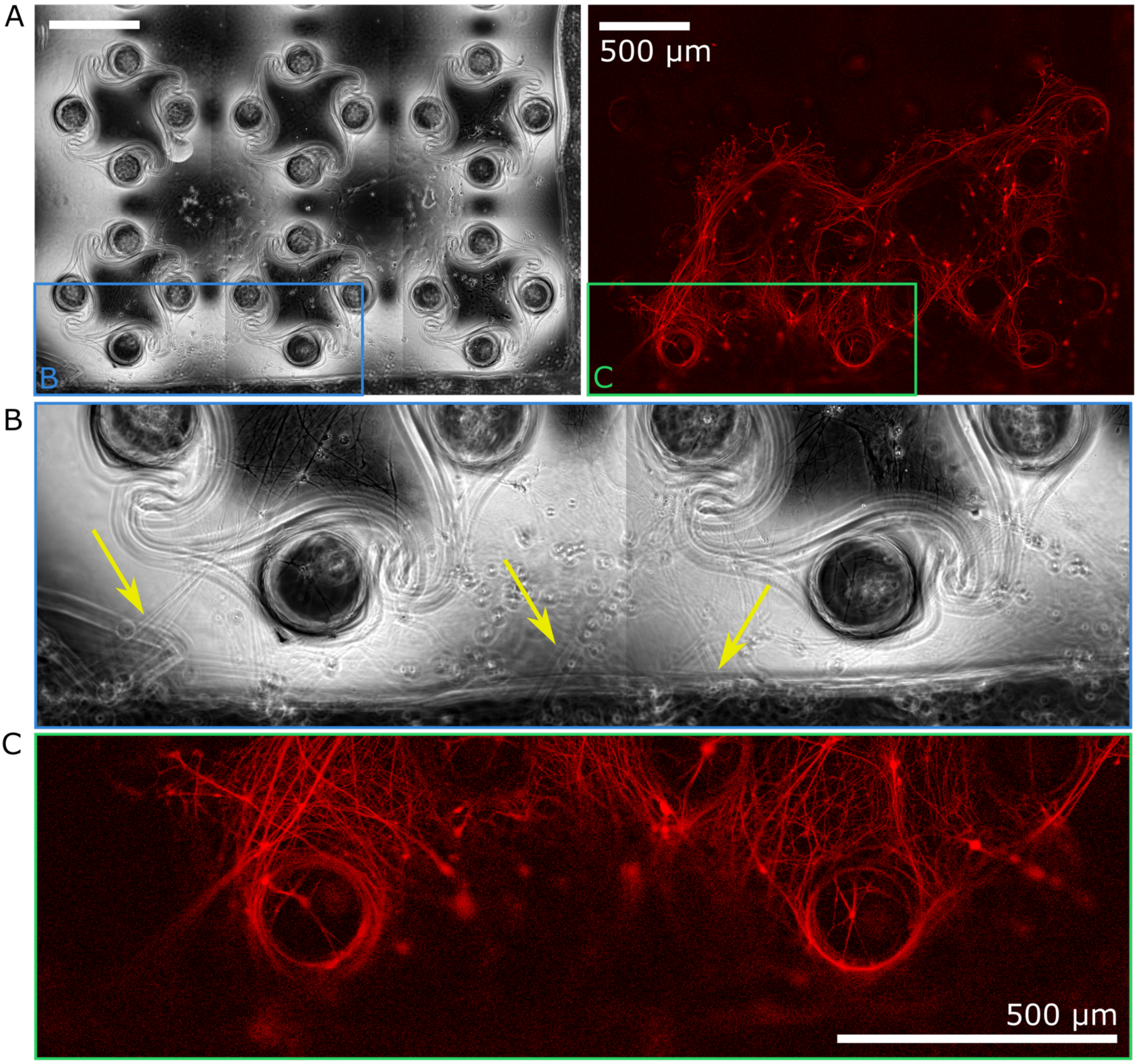


**Figure S3. Unwanted axonal growth on the top of a PFPA-PVP-coated PDMS microstructure**. **(A)** Overview of the six circuits presenting overgrowth, with RFP-expressing iNeurons. The blue and green rectangles are zoomed-in on B and C. **(B)** Phase-contrast zoom-in of the edge of the PDMS microstructure. Yellow arrows show axons connecting to the neurons located next to the PDMS microstructure. **(C)** Fluorescent zoom-in of the edge of the PDMS microstructure. These images were taken on a dissociated 5:1 iNeurons-glia sample.

#### 4 MORPHOLOGY OF CELLS GROWING NEXT TO THE PDMS

In dissociated samples, cell seeding relies on pipetting cells on top of the microstructures. While some of the cells land in the nodes of the PDMS microstructure, many cells land next to the PDMS microstructures. When imaging the circuits at different timepoints, images of the cells growing next to the PDMS microstructures were also taken. Fig. S4 shows such images, taken in a dissociated 5:1 iNeurons-glia sample at different timepoints.

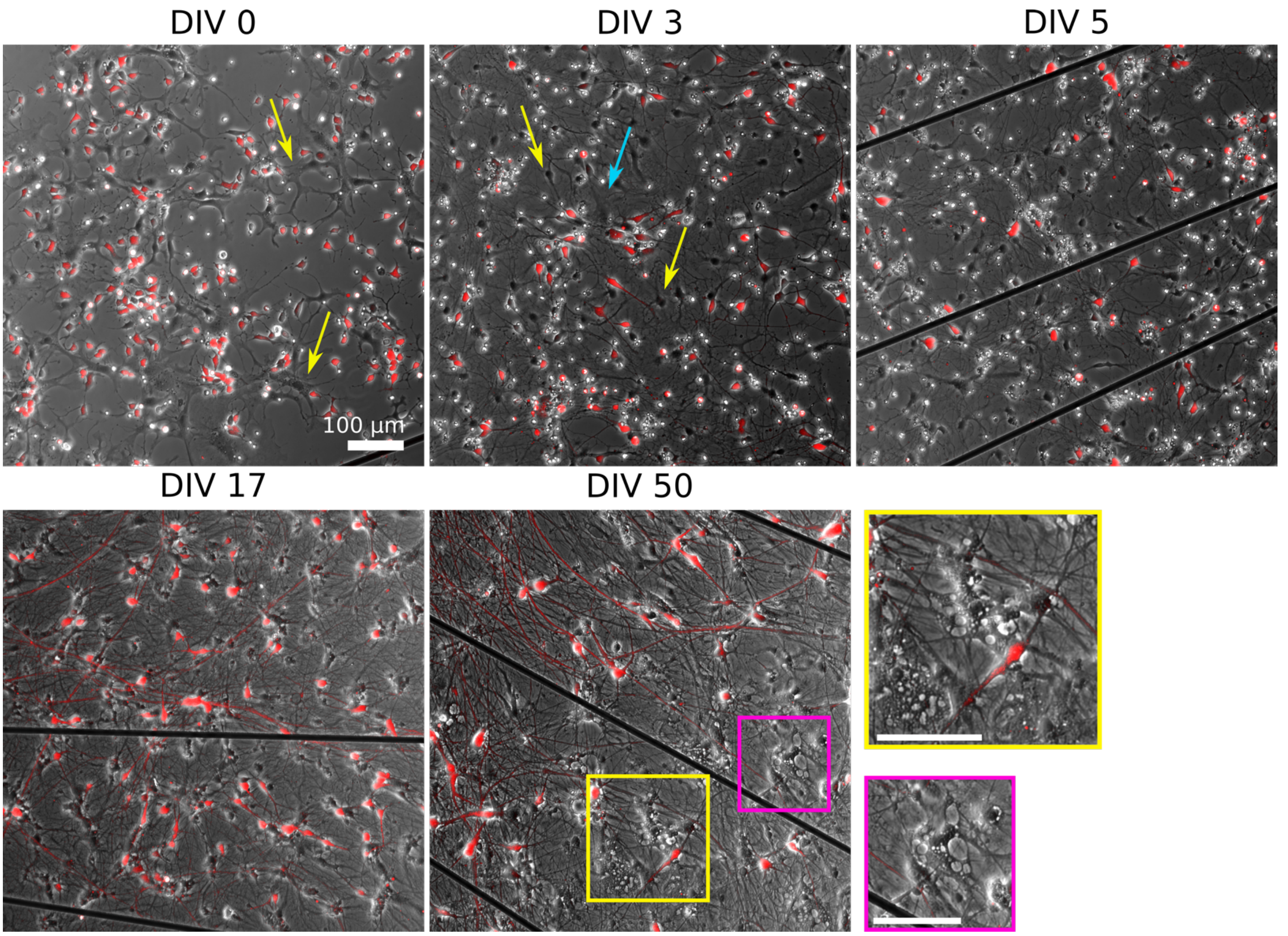


**Figure S4. Example images of cells outside of the PDMS microstructure at different timepoints.** RFP-expressing iNeurons are visible in red, while non-stained primary glial cells are visible in greyscale (phase-contrast image). These images were taken on a dissociated 5:1 iNeurons-glia sample.

These images show two elements that confirmed what was observed in the circuits in Fig. 5 and 6: first, dead cells disappear between DIV 5 and DIV 17, and second, the morphology of the glial cells change over time. Dead cells (likely neurons) appear as small, bright spots. A lot of them are visible on DIV 3 and DIV 5, but none are visible on DIV 17 and 50. The changes of morphology of glial cells, which are not stained, are easier to see on these phase-contrast images taken outside of the PDMS microstructures than on the phase-contrast images of the PDMS microstructures, where cells are confined together. At DIV 0, glial cells appear mostly as large, flat, spread-out cells with processes (yellow arrows). At DIV 3 and 5, most of the visible glial cells have a smaller and rounder cell body, more similar to the iNeurons (yellow arrows). These are likely astrocytes. Some spread out cells are still faintly visible (blue arrows). At DIV 50, some very large cells with vacuoles are visible (pink and yellow zoom-ins). These are likely microglia.

#### 5 MEA SAMPLES USED FOR ELECTROPHYSIOLOGY EXPERIMENTS

Fig. S5 shows images of a node of iNeurons without (A) and with (B) glial cells, taken at DIV 5, 17 and 50. In the iNeurons-only node, dead cells (small, bright, round spots) are present in the node in all three timepoints. In the glia-containig node, most dead cells disappear between DIV 5 and DIV 17. This is likely due to them getting phagocytosed by microglia.

Representative example circuits of iNeurons over time for the 3 different types of dissociated circuits. The red images show RFP-expressing iNeurons and the greyscale image is a phase-contrast view of the circuit, with the edges of the PDMS visible as a bright outline.

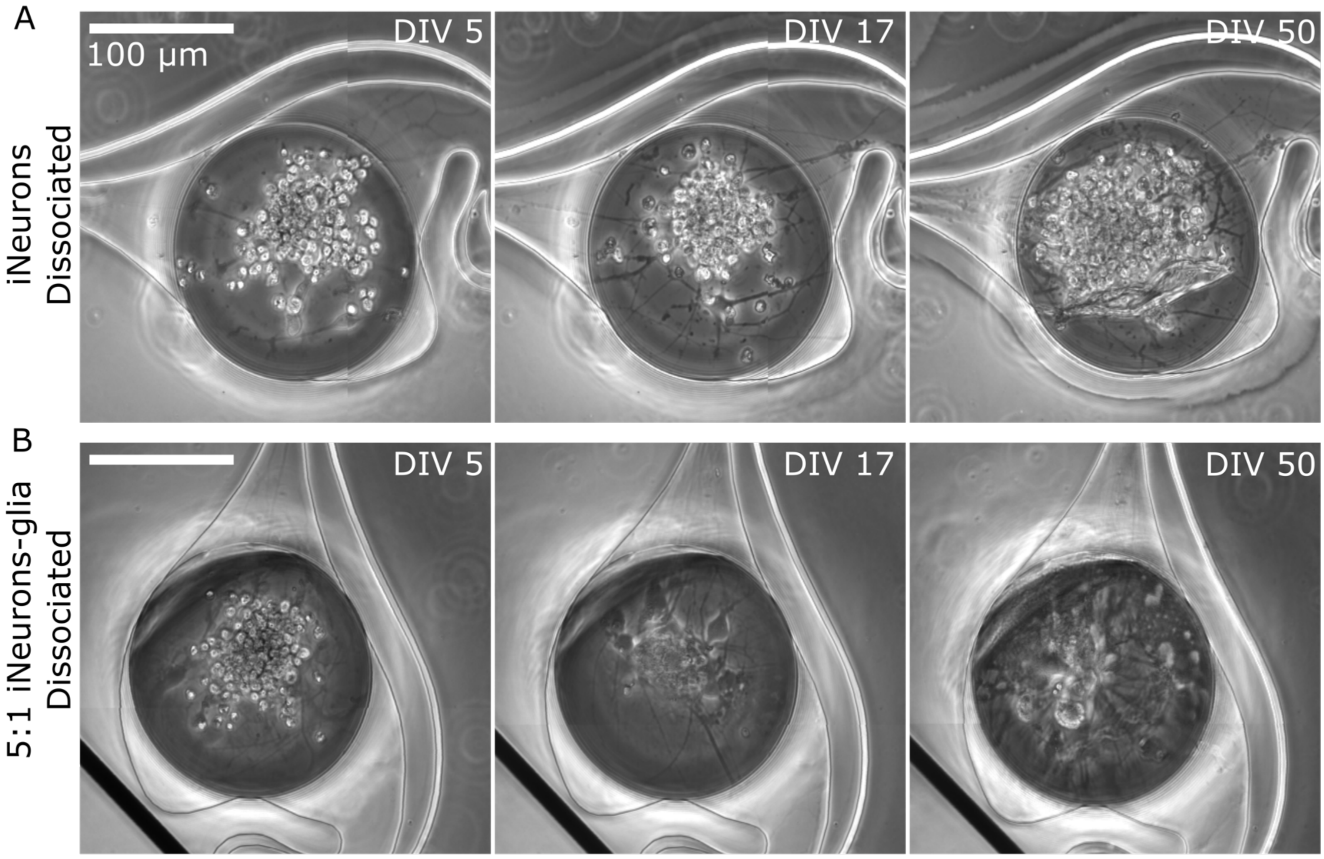


**Figure S5. Phase contrast images of nodes with dissociated iNeurons (A) and 5:1 iNeurons-glia (B)** Images were taken at DIV 5, 17 and 50. These are zoomed-in version of the nodes shown with blue arrows in Fig. 5A and 5B.

Fig. S6 shows the fluorescent images of the 18 MEAs with RFP-expressing neurons, taken at DIV 50. The PDMS lifting up can be observed on several images, for example in the first sample of Fig. S6D or on the first and third sample of Fig. S6F.

From the DIV 50 images shown in Fig. S6 and similar images taken at DIV 17, the percentage of non-empty nodes per MEA, *i.e.* the percentage of nodes with at least one live iNeuron, was calculated. Results can be seen in Fig. S7A. By grouping together the four nodes that belong to the same circuit, the percentage of circuits with four non-empty nodes (which we will refer to as “full” circuits) was also calculated (Fig. S7B). The highest percentage of full nodes and full circuits was in the iNeuron dissociated and spheroid samples (red). The glia containing samples generally had slightly more empty nodes, which resulted in fewer full circuits too. In all conditions but the 5:1 iNeuron-glia dissociated MEAs, the percentage of non-empty nodes slightly decreased between DIV 17 and 50. The largest decrease was in 5:1 iNeuron-glia samples, going from 94.4% of full nodes to 77.2%.

### 6 SPIKE TRAIN DIRECTIONALITY

As a complement to Fig. 9, Fig. S8 shows examples of spike trains at DIV 29, 43, and 49.

### 7 STIMULATION-INDUCED ACTIVITY

#### 7.1 250 ms-response upon stimulation

Fig. S9 shows the stimulation-induced electrical activity for the 6 different types of circuits. This plot is an extension of the data plotted in Fig. 10E, showing the full 250 ms rather than only the first 20 ms.

#### 7.2 Effect of extracellular $Mg^{2+}$ on the stimulation-induced spiking activity

Fig. S10 shows a zoom-in on the 20 ms of data shown in Fig. 13A and B.

#### 7.3 Comparing MFR and stimulation-induced activity

To test the effect of multiple medium changes and addition of small volumes of $H_2O$ and DMSO on the electrical activity of the network, we performed a reproducibility test with four sets of four recording sessions (A-D) (Fig. S11). Each recording session consisted in $1\times5$ min of spontaneous activity followed by $1\times10$ min of stimulation. Recording sessions were spaced by 20 min, during which samples were placed back into a 37°C incubator. Between recording session B and C, 5 µL of $H_2O$ and 5 µL of DMSO were added to the medium. After each set of four recordings, the medium was fully changed. After the third medium change, only two recording sessions were performed. This test was repeated on two samples: one sample with dissociated iNeurons and one sample with iNeuron spheroids. The MFR and mean number of induced spikes per stimulus were calculated for each repeat of spontaneous activity and stimulation-induced activity recordings, respectively. The coefficient of variation was calculated as the ratio of the standard deviation to the mean (over the fourteen measures shown on each plot).

Medium changes had an important effect on the MFR, with a coefficient of variation of 47.0 % for the dissociated iNeurons sample and of 43.1 % for the spheroid iNeurons sample. Over a single set of four measurements, the MFR usually increased, which might be linked to medium evaporation. The addition of a small volume of water and DMSO had little effect, or at least no more visible effect than the difference generally observed after waiting 20 min between two recording sessions. By contrast, the stimulation-induced electrical activity was much less impacted by medium change than the MFR, with a coefficient of variation of 10.0 % for the dissociated iNeurons sample and of 11.5 % for the spheroid iNeurons sample.

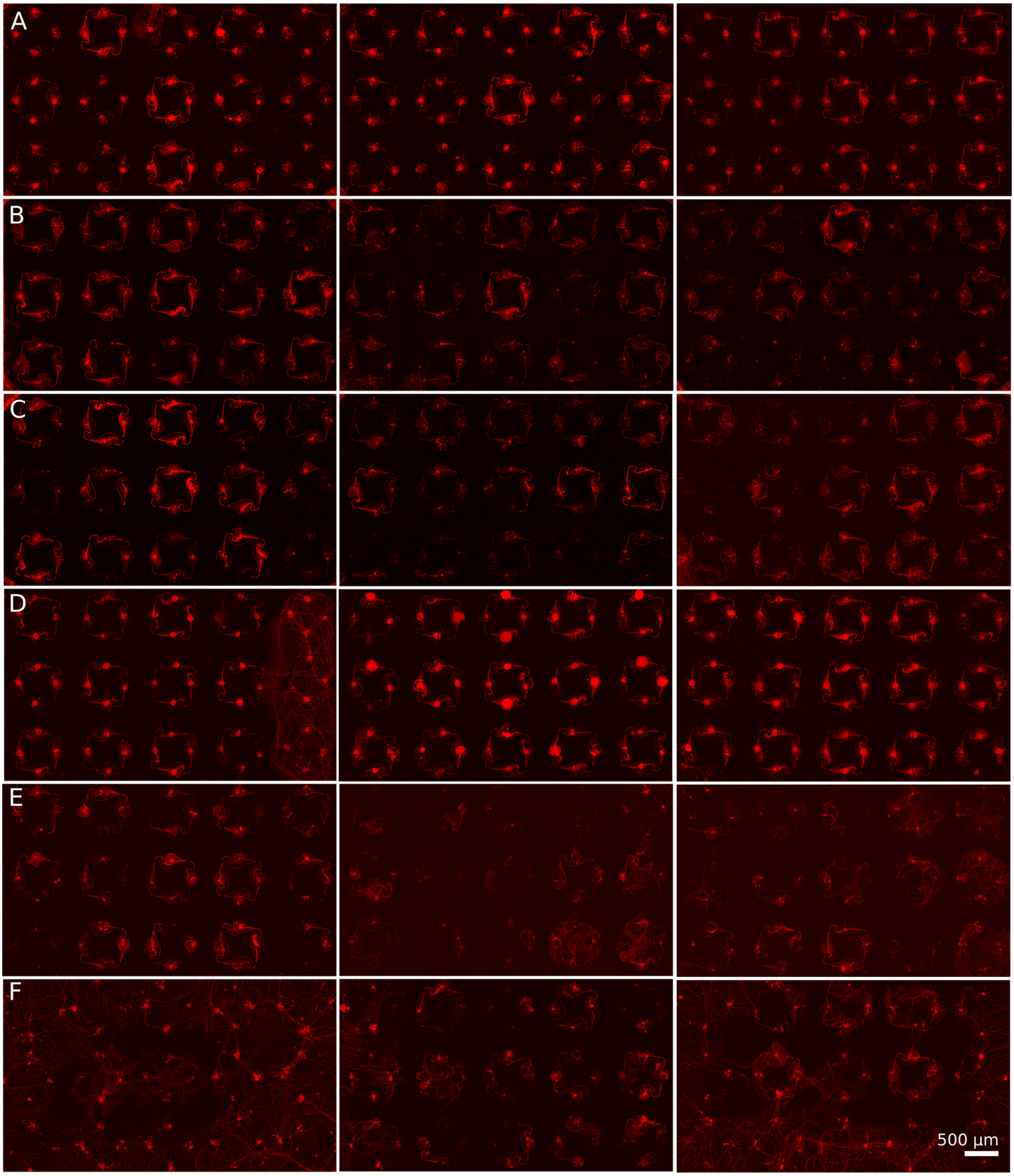


**Figure S6. Overview of the 3 MEAs of each conditions used for electrophysiology experiments.** Images were taken at DIV 50. RFP-expressing iNeurons are visible in red. **(A)** Dissociated iNeurons. **(B)** 5:1 iNeurons-glia (dissociated) **(C)** 2:1 iNeurons-glia (dissociated) **(D)** iNeurons spheroids **(E)** 5:1 iNeurons-glia spheroids **(F)** 2:1 iNeurons-glia spheroids

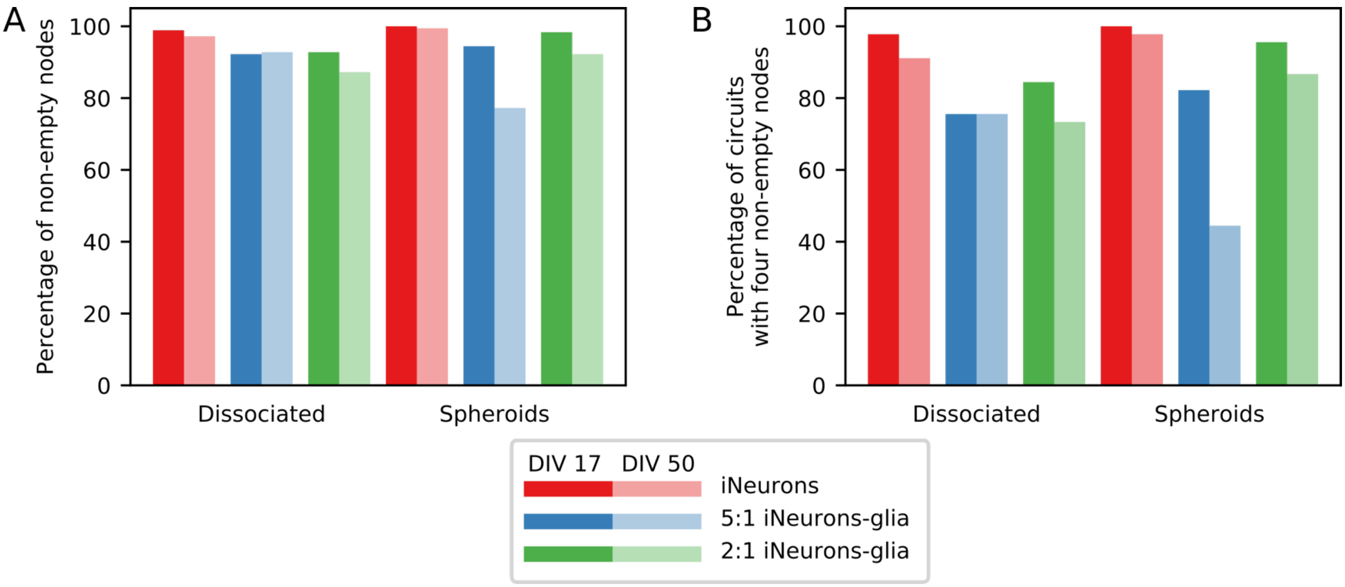


**Figure S7. Quantifying the number of nodes with at least one live iNeuron ("non-empty")** at DIV 17 and DIV 50 **(A)** Percentage of non-empty nodes per condition. **(B)** Percentage of circuits (four nodes) with all nodes full. Both of these plots are based on the same set of data, *i.e.* the count of empty nodes for each MEA at DIV 17 and DIV 50.

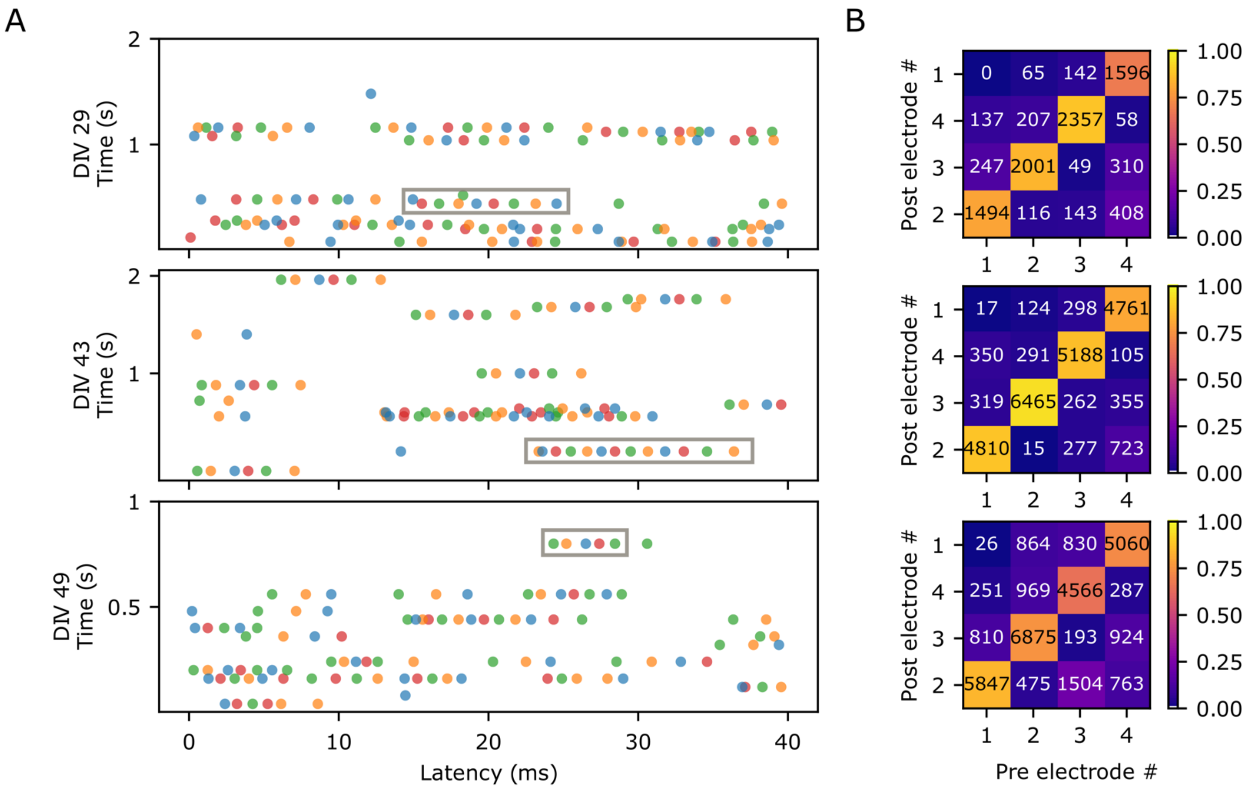


**Figure S8. Direction of action potential propagation in an example circuit (A)** Example spike trains, obtained from the spikes detected on the four electrodes of the circuit shown in Fig. 9. (B). Three different time points are shown: DIV 29 (2 sec), DIV 43 (2 sec) and DIV 49 (1 sec). The gray boxes highlight examples of consecutive spikes spaced by less than 5 ms and firing in the clockwise direction. **(B)** Frequency map of the occurrence of two consecutive spikes spaced by less than 5 ms, sorted by pre- and post-electrodes.

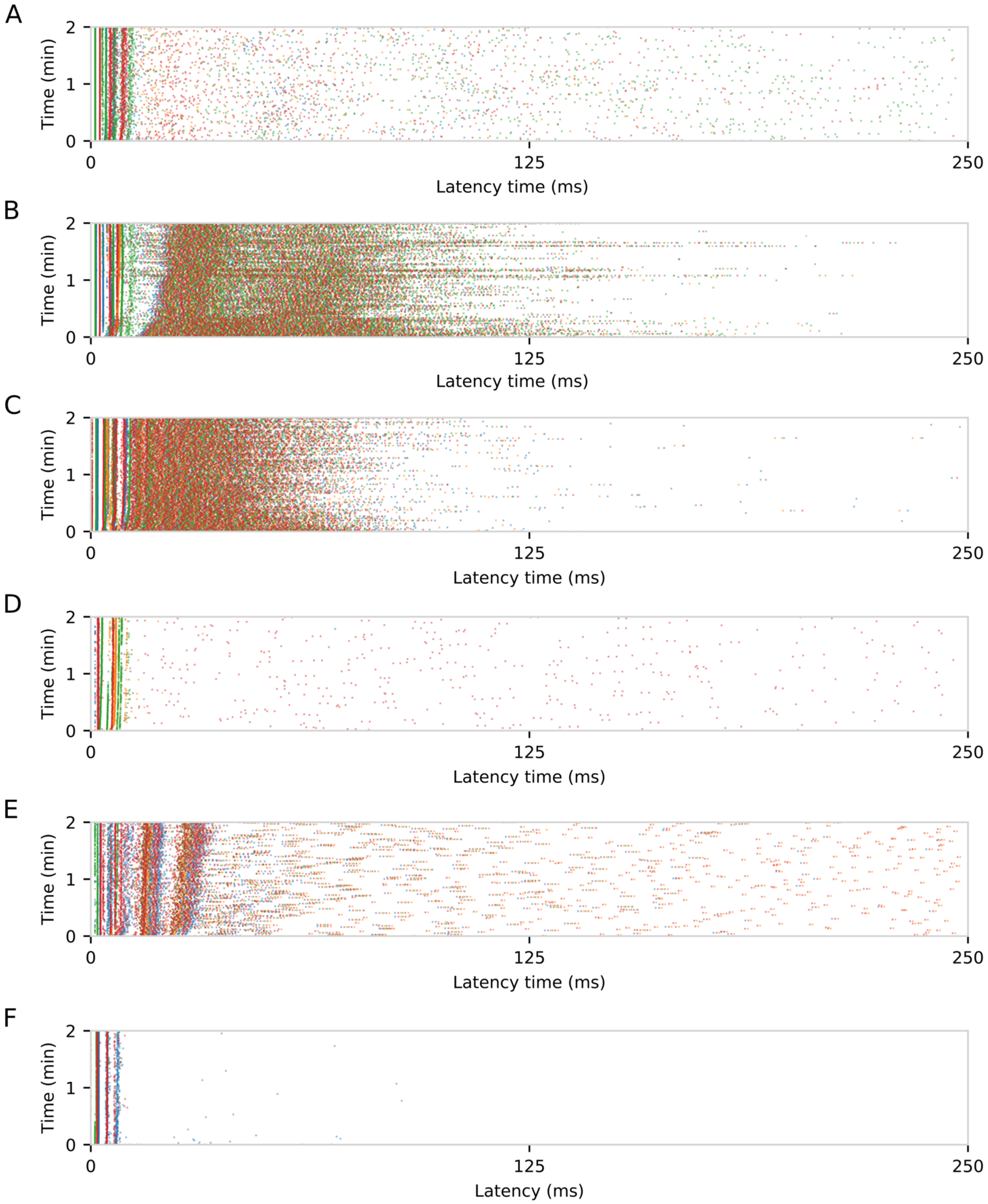


**Figure S9. Stimulation-induced electrical activity for the six different types of circuits**. This figure shows the extended data plotted in Fig. 10E, with the full 250-ms response rather than only the first 20 ms. **(A)** Dissociated iNeurons **(B)** Dissociated 2:1 iNeurons-glia **(C)** Dissociated 5:1 iNeurons-glia. **(D)** Spheroids of iNeurons. **(E)** Spheroids of 5:1 iNeurons-glia. **(F)** Spheroids of 2:1 iNeurons-glia.

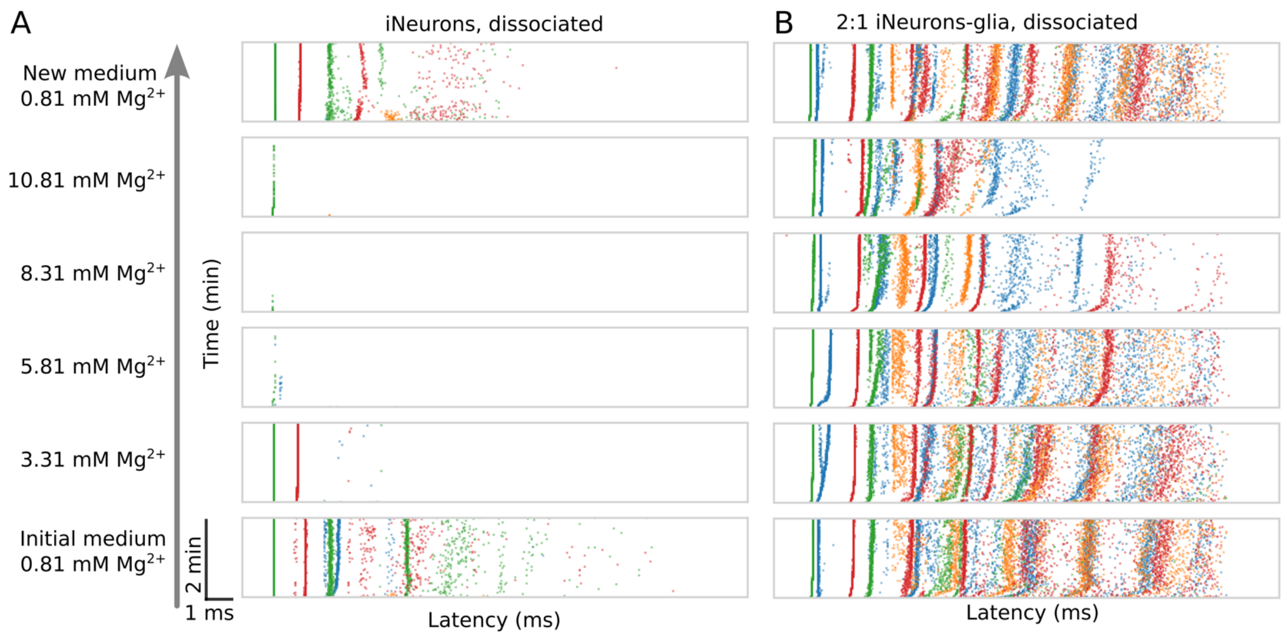


**Figure S10. Effect of extracellular $Mg^{2+}$ on the stimulation-induced electrical activity: the first 20 ms after stimulation**. This figure shows the a zoom-in of the data plotted in Fig. 13A and B **(A)** Example circuit of dissociated iNeurons. **(B)** Example circuit of dissociated 2-to-1 iNeurons-to-glia.

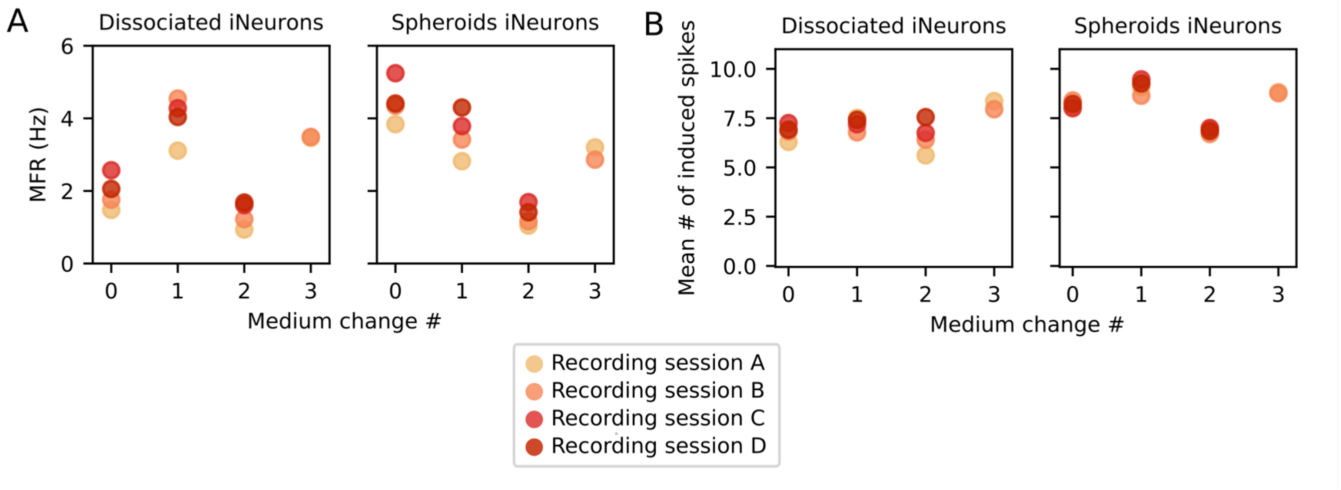


**Figure S11. Comparison of electrical activity metrics upon three sequential medium changes.** Between recording sessions B and C, 5 µL of $H_2O$ and 5 µL of DMSO were added to the medium to test if this impacted the electrical activity of the circuits. **(A)** Effect on the MFR. **(B)** Effect on the stimulation-induced activity.
